# Cryptochromes and UBP12/13 deubiquitinases antagonistically regulate DNA damage response in Arabidopsis

**DOI:** 10.1101/2023.01.15.524001

**Authors:** Yuzhao Hu, Daniele Rosado, Louise N. Lindbäck, Julie Micko, Ullas V. Pedmale

## Abstract

Cryptochromes (CRYs) are evolutionarily conserved blue-light receptors that evolved from bacterial photolyases that repair damaged DNA. Today, CRYs have lost their ability to repair damaged DNA; however, prior reports suggest that human CRYs can respond to DNA damage. Currently, the role of CRYs in the DNA damage response (DDR) is lacking, especially in plants. Therefore, we evaluated the role of plant CRYs in DDR along with UBP12/13 deubiquitinases, which interact with and regulate the CRY2 protein. We found that *cry1cry2* was hypersensitive, while *ubp12ubp13* was hyposensitive to UVC-induced DNA damage. Elevated UV-induced cyclobutane pyrimidine dimers (CPDs) and the lack of DNA repair protein RAD51 accumulation in *cry1cry2* plants indicate that CRYs are required for DNA repair. On the contrary, CPD levels diminished and RAD51 protein levels elevated in plants lacking UBP12 and UBP13, indicating their role in DDR repression. Temporal transcriptomic analysis revealed that DDR-induced transcriptional responses were subdued in *cry1cry2*, but elevated in *ubp12ubp13* compared to WT. Through transcriptional modeling of the timecourse transcriptome, we found that genes quickly induced by UVC (15 min) are targets of CAMTA 1-3 transcription factors, which we found are required for DDR. This transcriptional regulation seems, however, diminished in the *cry1cry2* mutant, indicating that CAMTAs are required for CRY2-mediated DDR. Furthermore, we observed enhanced CRY2-UBP13 interaction and formation of CRY2 nuclear speckles under UVC, suggesting that UVC activates CRY2 similarly to blue light. Together, our data reveal the temporal dynamics of the transcriptional events underlying UVC-induced genotoxicity and expand our knowledge of the role of CRY and UBP12/13 in DDR.

## INTRODUCTION

Genome integrity is essential for all living organisms, but it is especially important for plants because they are stationary and primarily grow post-embryonically. DNA damage can lead to the loss or alteration of essential genes, which can affect plant growth, development, and overall health^1^. Additionally, DNA damage can also lead to the production of abnormal proteins that can disrupt normal cellular processes or cause other negative effects^2^. Since plants rely on light as an energy source, they are inevitably exposed to DNA damage caused by both UV radiation in sunlight and the production of reactive oxygen species in chloroplasts due to excess light^3,4^. UV radiation is particularly damaging because it leads to the formation of cyclobutane pyrimidine dimer (CPD) in DNA, which inhibits DNA replication and RNA transcription^5^. Therefore, it is important for organisms to have effective DNA repair mechanisms to prevent and mitigate the negative consequences of DNA damage. Once such mechanism conserved in the eukaryotes is the DNA damage response (DDR), which generally encompasses three important aspects: 1) induction of DNA repair, 2) checkpoint response that halts the cell cycle, and 3) programmed cell death to eliminate cells with irreparable DNA damage^6^.

In plants and animals, similar DDR mechanisms exist, where the DNA repair and cell-cycle arrest are initiated by two kinases, ATAXIA TELANGIECTASIA-MUTATED AND RAD3-RELATED (ATR) and ATAXIA-TELANGIECTASIA MUTATED (ATM), which are activated by single-stranded DNA or DNA double-strand breaks (DSBs), respectively^7,8^. Upon activation, ATR and ATM target different downstream factors in animals and plants. In animals, ATR and ATM phosphorylate Checkpoint kinase 1 and 2 (Chk1 and Chk2), respectively^9^. ATR, ATM, Chk1, and Chk2 then phosphorylate and activate a master regulator of DDR, p53 transcription factor (TF), which induces the transcription of thousands of genes to orchestrate DDR^10^. In contrast, plant genomes lack orthologs of Chk1, Chk2 and p53^1^, instead, ATR and ATM activate a different TF, SUPPRESSOR OF GAMMA RADIATION 1 (SOG1), which is largely required for the induction of the transcriptional network of DDR^11^.

UV-induced CPD can be detected and repaired by the nucleotide excision repair (NER) pathway^12^. There are two NER pathways, the global genome NER (GG-NER), which scans the whole genome to detect CPD, and the transcription-coupled NER (TC-NER), which senses the CPD-induced stalling of RNA polymerase II during transcription^12^. Interestingly, both GG-NER and TC-NER require Cullin 4 (CUL4)-RING ubiquitin E3 ligase (CRL4): the former relies on CRL4^DNA DAMAGE BINDING 2 (DDB2)^ to recognize CPD, while the latter require CRL4^COCKAYNE SYNDROME A(CSA)^ to assemble the DNA repair machinery^13^. In plants, apart from NER, photolyases can harness energy from blue/UVA light to repair CPD without DNA excision^14^. Evolutionarily, photolyases are homologous to plant and animal cryptochromes (CRY)^15^. Plant CRYs perceive blue light to fine-tune growth and development^16^. Blue-light-activated CRYs dimerize and tetramerize to interact with downstream signaling partners^17^. Upon exposure to blue light, CRYs also form nuclear speckles^18,19^, where CRY2 carries out its function^20^. The photoactive CRYs are then ubiquitinated by two E3 ligase complexes, CONSTITUTIVE PHOTOMORPHOGENIC 1 (COP1) and SUPPRESSOR OF PHYA-105 (SPA) in complex with CRL4 (CRL4^COP1-SPA^), and Light Response BTB (LRB) in complex with CRL3 (CRL3^LRBs^) ^21,22^, and further degraded by 26S proteasomes^19,23^, desensitizing the CRY-mediated blue light signaling pathway^21,24^. Recently, we found that two deubiquitinases (DUBs), UBIQUITIN-SPECIFIC PROTEASE 12 (UBP12) and UBP13 (UBP12/13), interact with CRY2 in a blue light-dependent manner and stabilize COP1^25^, which contributes to the blue light-specific degradation of CRY2 mediated by COP1^25^.

Ubiquitination marks are important for both plant and animal DDR. For example, SOG1 in plants is stabilized through ubiquitination by DNA DAMAGE RESPONSE MUTANTS 1 (DDRM1), contributing to the homologous recombination (HR) repair upon DNA damage^26^. Moreover, animal p53 is ubiquitinated by the E3 ligase MURINE DOUBLE MINUTE 2 (Mdm2), leading to the degradation of p53 and the depletion of p53 from the nucleus^27^. Therefore, DUBs that removes the ubiquitination marks have been identified as important regulators of DDR, especially in animals. For example, the UBP12/13 homolog in animals, ubiquitin-specific protease 7 (USP7), deubiquitinates p53 and Chk1 to stabilize them^28^. In addition, USP3 deubiquitinates histone H2A and H2AX, negatively regulating the recruitment of DNA damage repair proteins^29^. However, despite the role of ubiquitination in the DDR being largely conserved in plants, how plant DUBs are regulating the DDR remains largely unexplored.

Present-day CRYs have lost their enzymatic activity to repair pyrimidine dimers directly^30^, but continue to bind preferentially to CPD-containing DNA and regulate DDR in mammals^31,32^. Evidence in *Arabidopsis* suggests that CRYs affect the activity of the UVB receptor UV RESISTANCE LOCUS 8 (UVR8) in a blue-light dependent manner and, therefore, are involved in UVB tolerance under natural light conditions^33–35^.

However, the role of plant CRYs in the DDR is not well understood. In this study, we demonstrate that CRYs act as positive regulators of DDR in plants when exposed to UVC, promoting the repair of UV-induced CPDs and the expression of DDR-related genes. Additionally, we show two known CRY2 regulators, UBP12 and UBP13 DUBs, counteract the activity of CRYs, thus revealing the important role of deubiquitinases in DDR in plants. Through transcriptomic analysis of CRY and UBP12/13 mutants during DDR, we identify CALMODULIN-BINDING TRANSCRIPTION ACTIVATORs (CAMTAs) as novel regulators of DDR that potentially act downstream of the CRY-UBP12/13 module. Furthermore, we also find that UVC enhances CRY2-UBP13 interaction, resulting in the degradation of CRY2. Furthermore, we observed the formation of punctate nuclear bodies (photobodies) of CRY2 upon UVC exposure. In summary, we reveal that the CRYs-UBP12/13 module is harnessed by plants to not only optimize growth in accord with visible light signals, but also establish resistance against UV-caused DNA damage.

## RESULTS

### CRY1/2 promote, whereas UBP12/13 inhibit resistance against DNA damage

To elucidate the role of CRY1 and CRY2 (CRY1/2) and UBP12/13 in the DDR, we examined the phenotype of *Arabidopsis* single and double CRY1 and CRY2 mutants along with UBP13 overexpressing seedlings (*UBP13oe*) and the double mutant *ubp12ubp13*, as UBP12 and UBP13 are genetically redundant^25,36^. To induce DDR, we employed UVC, which is a known genotoxin^37^. We treated 4-day-old light-grown seedlings with 0, 5500, 8000 J/m^2^ UVC, then grew them back under light for 6 days before inspecting their phenotype (Figure S1A). At 5500 and 8000 J/m^2^ UVC, *cry1, cry2, cry1cry2* and *UBP13oe* showed pale cotyledons, while *ubp12ubp13* and wild-type (WT) did not (Figures 1A and S1B). To assess plant growth and survival after DNA damage, we measured seedling weight after UVC treatment. The *cry1, cry2, cry1cry2* mutants and *UBP13oe* line had lower weights (Figures 1B and S1C), while *ubp12ubp13* had a ~1.5 times higher weight than the WT after UVC (Figures 1B). These results suggest that CRY1/2 positively regulate while UBP12/13 negatively regulate DDR. UBP13 interacts with CRY2 and regulates its blue light signaling pathway^25^. Therefore, we examined whether UBP12/13 also act in the same genetic pathway as CRY1/2 during DDR. We examined *cry1cry2;UBP13oe* seedlings upon DNA damage and found that *cry1cry2;UBP13oe* exhibited pale cotyledons and had a lower fresh weight than WT (Figures 1A and 1B), phenocopying *cry1cry2* mutants (Figures 1A and 1B), indicating that UBP13 functions in the same genetic pathway as CRY1/2 to regulate DDR.

**Figure 1.**
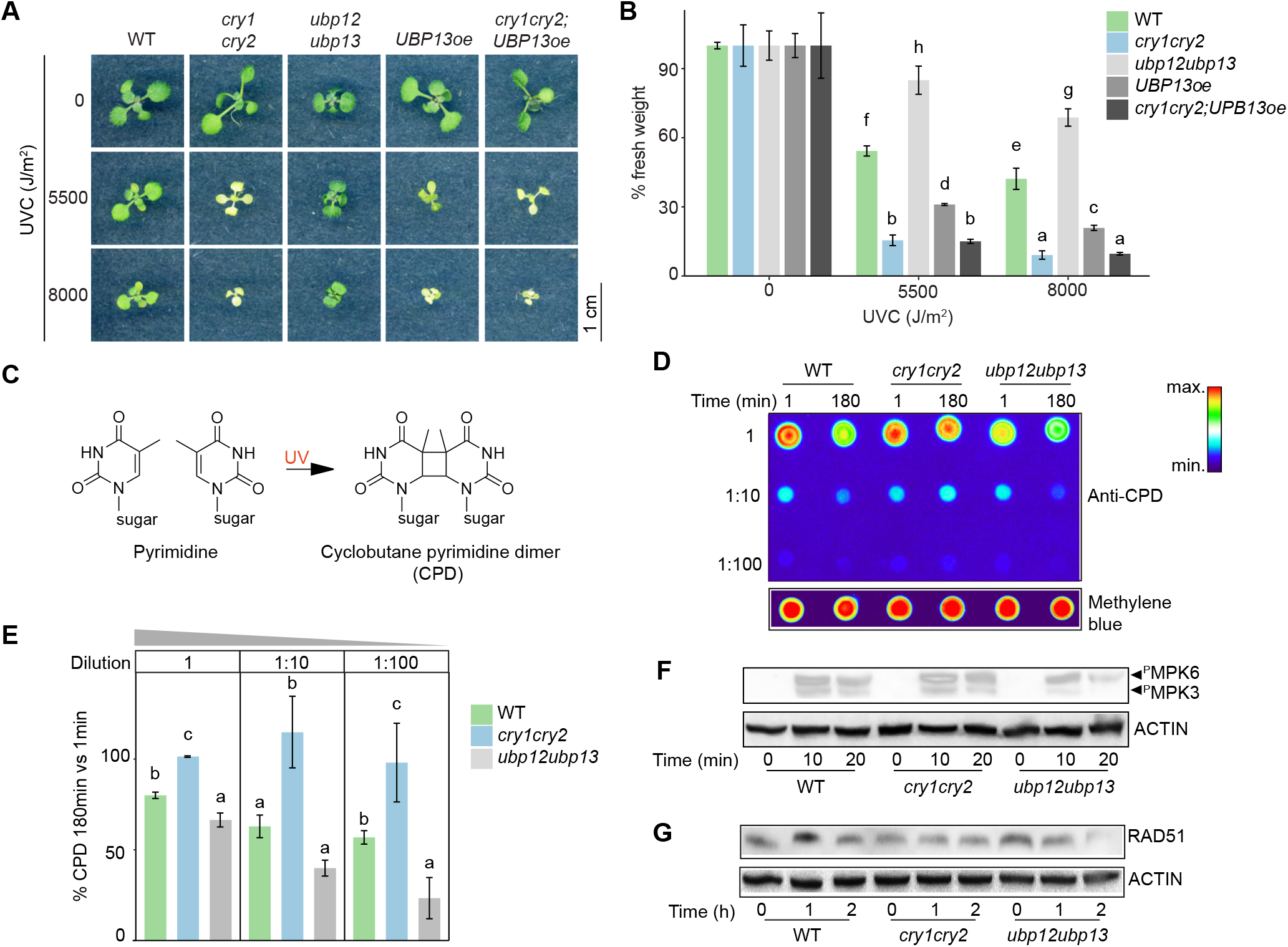
*Arabidopsis* plants deficient in CRY1 and CRY2 are susceptible to UVC-induced DNA damage, while mutants of *UBP12* and *UBP13* are not. (A) Phenotype of representative 10-day-old seedlings of the indicated genotypes treated with indicated intensities of UVC. 4-day-old light-grown seedlings were treated with UVC and then returned to white light for 6 days before examination of the phenotype. (B) Fresh weight of 10-day-old seedlings of the indicated genotypes treated with indicated UVC doses as described in (A). Fresh weight was normalized to the untreated (0 J/m^2^) samples of the same genotype. (C) Schematic diagram illustrating the UV-induced formation of a CPD from two adjacent pyrimidines (two thymine bases shown as an example). (D) Representative dot blot showing CPD levels on the genomic DNA from the indicated genotypes. After treating 5-day-old seedlings with 6000 J/m^2^ UVC, genomic DNA was extracted after 1 or 180 minutes and serially diluted and CPD was detected using an anti-CPD antibody by immunoblotting. Serial dilutions of the genomic DNA were blotted: 1, 1:10, and 1:100. The immunoblot has been pseudo-colored to reflect the difference in intensities of CPD levels. Methylene blue staining shows equal loading of genomic DNA. (E) Quantification of CPD levels using replicates of the dot blot shown in (D). The percentage of CPD was derived by normalizing the CPD level at 180 min to 1 min of the same genotype. One-way analysis of variance (ANOVA) analysis was separately performed for each serial dilution. (F, G) Immunoblot analysis of phosphorylated MPK6 and MPK3 (F) and RAD51 (G) in indicated genotypes at indicated time points after UVC exposure. 5-day-old seedlings were untreated (0 min) or treated with 6000 J/m^2^ UVC and collected after 10 or 20 min in (F) and 1 or 2 h in (G). ACTIN was used to normalize the amount of total protein. For (B) and (E), different letters indicate *p*<0.05 for one-way ANOVA analysis followed by Fisher’s least significant difference (LSD) posthoc test. Data show means ± standard deviation (SD), n = 3 independent replicates.

Next, to explore whether CRYs are involved in resistance against other genotoxins, we used zeocin, which induces DSBs^38^. *KU70* is required for DSB repair^39^, therefore, we used the *ku70* mutant as a positive control. We treated 4-day-old WT, *ku70, cry1cry2* seedlings with 0, 4, and 8 μM zeocin for 8 days. As expected, *ku70* was hypersensitive to zeocin as it had a lower fresh weight than WT (Figure S1D). However, the fresh weight of *cry1cry2* and WT was similar after zeocin treatment (Figure S1D), indicating that CRYs are not required to resist DSB. Therefore, we focused on the UVC-induced DDR for the scope of our study. Our combined results suggest that CRY1 and CRY2 both play a positive role in DDR. Furthermore, UBP12 and UBP13 function in the same genetic pathway as CRY1/2 to negatively regulate DDR.

### Loss of CRY1 and CRY2 leads to higher CPD accumulation and lower DNA repair

Absorption of UV results predominately in CPD-type DNA damage, where cytosine (C) to thymine (T) or CC to TT mutations occur^5^ (Figure 1C). Changes in CPD levels can be used to track the progress of DNA damage repair after UVC exposure^40^. Therefore, to examine whether DNA damage repair is misregulated in *cry1cry2* and *ubp12ubp13*, we measured CPD levels in these mutants by dot-blot analysis after UVC exposure. 5-day-old light-grown plants were treated with UVC (6000 J/m^2^) and recovered in light for 1 min and 180 min. Genomic DNA was then extracted and dot-blotted at different serial dilutions to increase the dynamic range of detection. Using an anti-CPD antibody, we detected CPD in WT, *cry1cry2*, and *ubp12ubp13* 1 min after UVC exposure (Figures 1D and S1E), indicating accumulation of DNA damage in all three genotypes. In WT, the accumulated CPD decreased at 180 min suggesting CPD repair, as expected, while *ubp12ubp13* exhibited even lower CPD levels at 180 min (Figures 1D, E and S1E), suggesting enhanced CPD repair. Importantly, CPD levels in *cry1cry2* remained mostly unchanged 180 min after UVC exposure (Figures 1D, E and S1E), indicating impaired CPD repair in this mutant.

UVC and UVB stresses induce the phosphorylation and consequent activation of MAP KINASE 3 (MPK3) and 6 (MPK6)^41,42^, which are gradually dephosphorylated as plants recover from the stress^42^. Also, in this context, deficiencies in CPD repair are related to the hyper-phosphorylation of the two kinases^43^. Similarly, mutants with sustained levels of phosphorylated MPK3 (MPK3^P^) and MPK6^P^ following UVB radiation are hypersensitive to this genotoxic treatment^42^. Since we observed differences in sensitivity to UVC-induced DNA damage and CPD repair rates, we wondered if MPK3 and MPK6 were differentially phosphorylated in *cry1cry2* and *ubp12ubp13* mutants relative to WT. For this, we extracted proteins from 5-day-old seedlings that were either untreated (0 min) or treated by UVC and collected after 10 min or 20 min of recovery. We detected the phosphorylated forms of the two kinases in an immunoblot using an anti-MPK3^P^ and -MPK6^P^ antibody^42^. In all three genotypes, MPK3^P^ and -MPK6^P^ were only detected after UVC treatment and decreased after 20 min of recovery (Figure 1F). We then quantified the levels of MPK6^P^, which is the most abundant of the two detected MPKs (Figure 1F). Compared to WT, levels of MPK6^P^ were increased in the hypersensitive *cry1cry2* and decreased in the resistant *ubp12ubp13* after UVC (Figures 1F and S1F). Therefore, MPK6^P^ dephosphorylation during recovery was delayed in *cry1cry2* mutant and accelerated in *ubp12ubp13*, correlating with CPD levels in these genotypes (Figures 1D, 1E and S1E), which might explain their different sensitivities to UVC treatment (Figures 1A and 1B).

Apart from CPDs, UV also induces DSBs, the repair of which requires BREAST CANCER GENE 1 (BRCA1) and RAD51^43^. Moreover, DNA damage induces the expression of *BRCA1* and *RAD51*^44^. Therefore, to check for the DNA repair activity in WT, *cry1cry2* and *ubp12ubp13*, we measured the expression of *BRCA1* and *RAD51* in these three genotypes 1.5 h after UVC by reverse transcription-quantitative polymerase chain reaction (RT-qPCR). In WT, *BRCA1* and *RAD51* were induced by UVC compared to untreated samples (Figure S1G), suggesting normal DNA repair activity. In contrast, *BRCA1* and *RAD51* were not induced in *cry1cry2* upon UVC (Figure S1G), suggesting impaired DNA repair activity. Moreover, *BRCA1* and *RAD51* gene expression in *ubp12ubp13* was already higher than WT in untreated samples, then diminished after UVC treatment (Figure S1G), suggesting that *ubp12ubp13* had increased DNA repair activity. Because RAD51 protein is directly involved in DNA damage repair and DNA damage induces the accumulation of RAD51 protein^45–47^, we next measured the accumulation of RAD51 protein after UVC to corroborate the qPCR assays. We treated 5-day-old seedlings with UVC (6000 J/m^2^) and allowed them to recover for either 1 or 2 h before total proteins were extracted for immunoblot analysis using an anti-RAD51 antibody. In WT, RAD51 levels increased at 1 h after UVC exposure when compared with the untreated samples (0 h) and then recovered to basal levels at 2 h indicating normal DNA repair activity (Figures 1G and S1H). In contrast, induction of RAD51 was not observed in *cry1cry2* at either 1 h or 2 h after UVC exposure indicating the absence of DNA repair activity (Figures 1G and S1H). Interestingly, RAD51 levels were already higher in *ubp12ubp13* untreated seedlings when compared to WT and *cry1cry2*, but its levels diminished at 1 h and 2 h after UVC (Figures 1G and S1H), suggesting enhanced DNA repair activity. Altogether, these results indicate that *cry1cry2* has impaired while *ubp12ubp13* has enhanced DNA repair activity under UVC. Collectively, these findings reinforce that CRYs positively mediate DDR, while UBP12/13 negatively regulate DDR.

### CRYs promote while UBP12/13 inhibit the transcriptional response to UVC

To obtain further insights into how genetic losses of CRYs and UBP12/13 affect the transcriptional response to DNA damage, we analyzed a time-course transcriptome by RNA-sequencing (RNA-seq) in WT, *cry1cry2*, and *ubp12ubp13* after UVC. First, to select optimal sampling time points for the RNA-seq, we did an exploratory RT-qPCR assay in WT at 0, 15, 30, 60, 120 and 180 min after UVC. We then analyzed four known UV-responsive genes, *BRCA1, RAD51, PHOTOLYASE 1 (PHR1)* and *CINNAMATE-4-HYDROXYLASE* (*C4H*)^37,48,49^ (Figure S2A). We found that *BRCA1* was strongly induced early in our time course, 15 min after the UVC treatment (Figure S2A). *PHR1* expression peaked at 60 min (Figure S2A), while *BRCA1, RAD51* and *C4H* peaked at 180 min after UVC (Figure S2A). Since these three time points (15, 60 and 180 min) seemed to cover early as well as peak induction of our selected marker genes (Figure S2A), they were chosen for the RNA-seq experiment.

5-day-old seedlings of WT, *cry1cry2* and *ubp12ubp13* were harvested before (0 min) and 15, 60 and 180 min after UVC treatment (Figure 2A). Biological replicates in the RNA-seq are highly similar as evidenced by the high Pearson’s correlation coefficient (0.92 to 1) (Figure S2B) and the close proximity of replicates in the principal component analysis (PCA) plot (Figure S2C). To explore the pathways by which CRYs and UBP12/13 regulate the transcriptional response to DNA damage, we obtained differentially expressed genes (DEGs) from the RNA-seq. We identified DEGs (defined as false discovery rate-adjusted *p* value (*q* value) < 0.05) for each genotype by comparing the gene expression at 15 min, 60 min and 180 min to 0 min, respectively. DEGs in *cry1cry2* and *ubp12ubp13* relative to WT were obtained by comparing the gene expression in the corresponding mutant to WT at all four time points.

**Figure 2.**
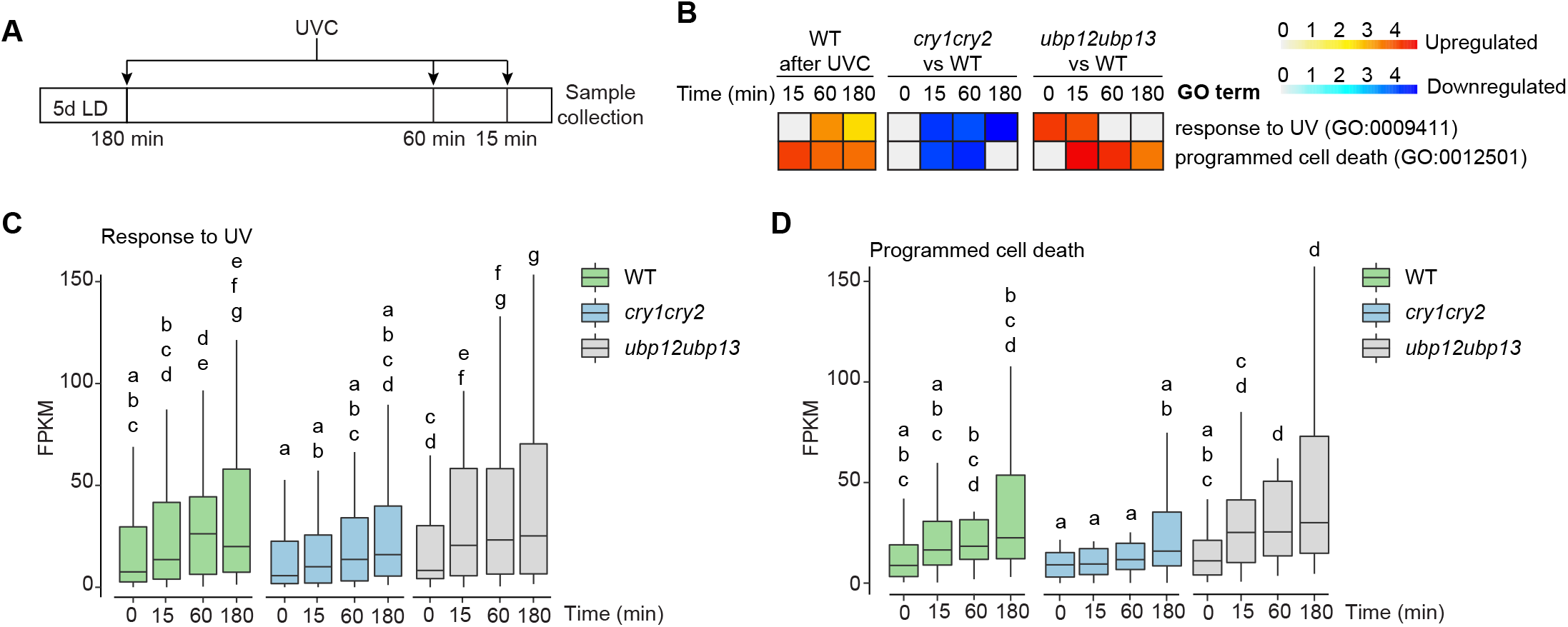
CRYs promote while UBP12 and UBP13 inhibit DNA damage-induced stress response. (A) Schematic diagram illustrating the UVC treatment for the RNA-seq samples. 5-day-old light-grown seedlings were treated with 6000 J/m^2^ of UVC and collected after 0, 15, 60, and 180 min of recovery time in white light. (B) Heatmap of the GO terms “response to UV” and “programmed cell death” in the upregulated genes in WT, downregulated genes in *cry1cry2* relative to WT and upregulated genes in *ubp12ubp13* relative to WT at indicated time points after UVC. The fold enrichment of non-statistically significant GO terms (false discovery rate ≥0.05) was manually set to 0. (C, D) Boxplot showing the expression levels (fragments per kilobase million, FPKM) of genes in the GO term “response to UV” (C) and “programmed cell death” (D) in WT, *cry1cry2* and *ubp12ubp13.* Only genes downregulated in *cry1cry2* relative to WT are shown. Different letters indicate *p*<0.05 for two-way ANOVA analysis followed by Fisher’s LSD posthoc test.

We then used these DEG lists to perform gene ontology (GO) enrichment analysis (Table S1). The top 10 GO terms enriched in the upregulated genes in WT at 15, 60 and 180 min included many stress-related GO terms (Figure S2D, Table S1), consistent with known studies reporting that UVC can induce stress responses^50^. Importantly, in *cry1cry2* and *ubp12ubp13* these stress-related GO terms were enriched in the downregulated and upregulated genes relative to WT, respectively (Figure S2D), suggesting that the transcriptomic response to UVC-induced DNA damage is less prominent in *cry1cry2* and enhanced in *ubp12ubp13.* Next, we found that the GO term “response to UV” was significantly enriched in the genes upregulated in WT (Figure 2B, Table S1), as expected. Importantly, this GO term was enriched in genes downregulated in *cry1cry2* and upregulated in *ubp12ubp13* relative to the WT (Figure 2B, Table S1). The expression of genes in the GO term “response to UV” was not significantly changed in *cry1cry2* or *ubp12ubp13* relative to WT at 0 min, but was lower in *cry1cry2* and higher in *ubp12ubp13* compared to WT at 60 min (Figure 2C, Table S2), suggesting that, after UVC treatment, genes in the known UV-responsive pathways are less induced in *cry1cry2* and more induced in *ubp12ubp13* compared to WT. Upon UV-induced DNA damage, DDR can trigger programmed cell death to protect genome integrity^51^. Accordingly, the GO term “programmed cell death” is highly enriched in the upregulated genes in WT (Figure 2B). Moreover, this GO term is enriched in the downregulated genes in *cry1cry2* and upregulated genes in *ubp12ubp13* relative to WT (Figure 2B). The expression of genes involved in the GO term “programmed cell death” also showed less induction in *cry1cry2* at 60 min compared to WT (Figure 2D, Table S2), suggesting that the UVC-induced programmed cell death may be diminished in *cry1cry2*, possibly contributing to the hypersensitive phenotype of *cry1cry2* (Figures 1A and 1B). Altogether, these results suggest that DNA damage-induced transcriptional responses are weaker in *cry1cry2* and stronger in *ubp12ubp13* compared to WT.

### CAMTAs mediate DDR

To gain insights into how the DEGs in WT are coordinately regulated during the UVC-induced DDR, we performed a Dynamic Regulatory Events Miner (DREM) analysis to generate a model of the underlying gene regulatory network^52,53^. DREM analysis takes time-course gene expression data as input, assigns genes into different groups based on similar expression patterns and predicts which transcription factors might be responsible for modulating the expression of each group of genes^52,53^. We first analyzed the expression of the DEGs in the WT along the time course after UVC and uncovered seven groups of co-expressed genes (W1-7) in the DREM model (Figure 3A). In the WT DREM model, SOG1, WRKYs and CAMTAs were predicted to regulate the induction of gene expression upon UVC treatment (Figure 3A). Since CAMTAs were predicted to regulate the most upregulated branch (consisting of paths W1, 2 and 7) for the early gene expression change at 15 min (Figure 3A), we next focused on studying the role of CAMTAs in the UVC-induced DDR. First, to corroborate the DREM prediction, we performed a *de novo* motif search in the promoters of the genes within the W1, 2 and 7 paths (Figure 3A, Table S2)^54^, and found a highly enriched “CGCGTT” motif (Figure 3B), which is a known CAMTA-binding DNA element, the rapid stress response element^55^, suggesting the CAMTAs can bind to the promoters of genes in the W1, 2, and 7 paths. Furthermore, when all DEGs were considered, *de novo* motif search identified CAMTA-binding motifs enriched in the promoters of only the upregulated but not downregulated genes in WT after UVC treatment (Figures S3A and S3B). These results suggest that CAMTAs might be required for the induction of DEGs in WT during DDR. CAMTA 1, 2 and 3 are involved in various abiotic and biotic stress responses^56^, but their possible role in DDR remains unexplored. To address this, we treated *camta1camta2camta3* (hereafter *camta123*) triple mutant with UVC. Similar to *cry1cry2*, the *camta123* mutant had pale cotyledons and a lower fresh weight than WT after UVC treatment (Figures 3C and 3D), suggesting that CAMTAs are indeed required for DDR.

**Figure 3.**
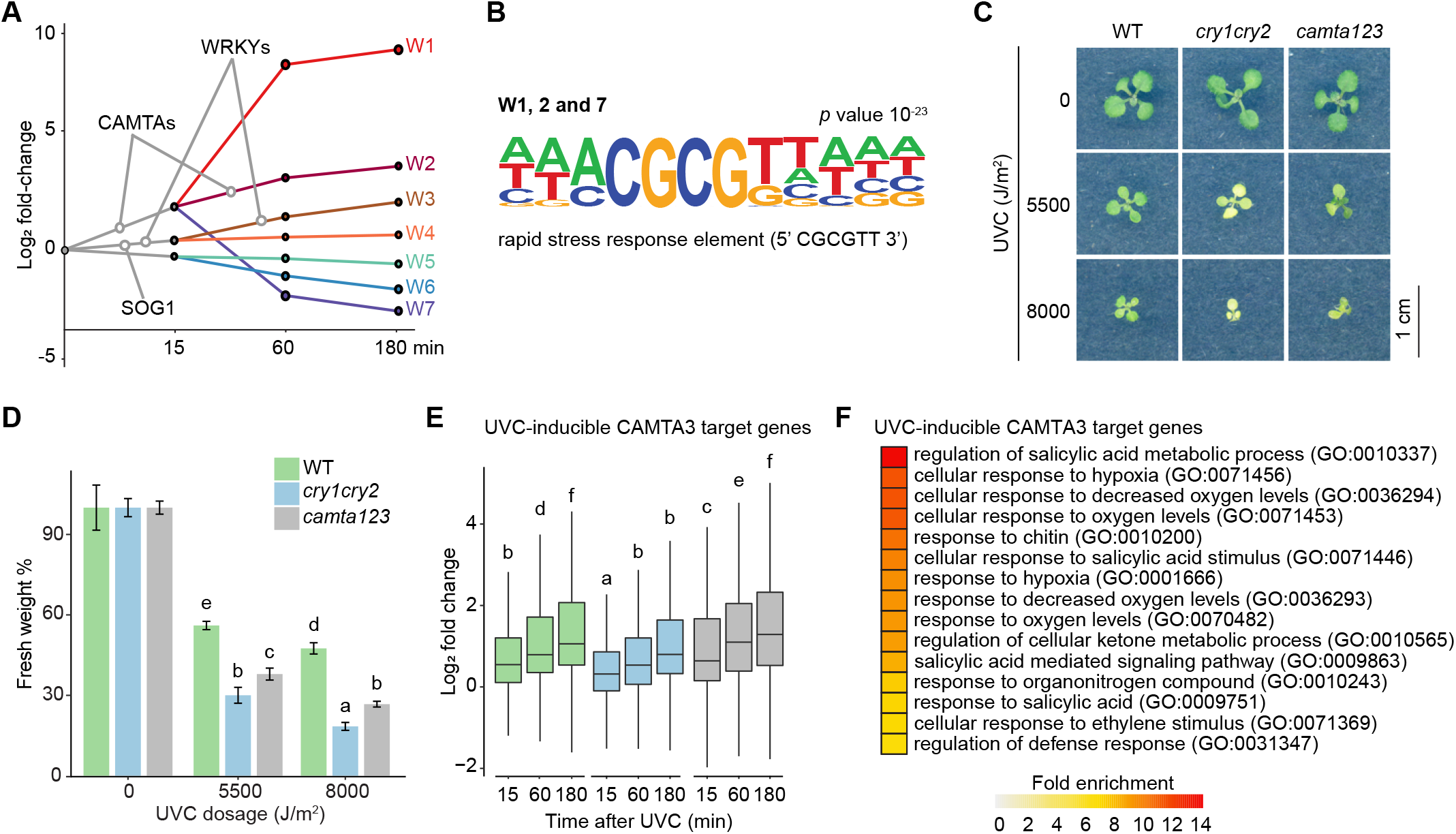
CAMTAs are required for DDR. (A) Groups of co-expressed DEGs in WT after UVC. 7 co-expressed groups of genes (W1-7) were identified. CAMTAs, SOG1 and WRKYs TFs were predicted to regulate the indicated groups. (B) Consensus sequence of the top *cis*-regulatory motif found in the promoters of W1, 2 and 7 genes shown in (A). (C) Phenotype of representative 10-day-old seedlings of the indicated genotypes treated with indicated UVC doses. 4-day-old light-grown seedlings were treated with UVC and then returned to white light for 6 days before examination of the phenotype. (D) Fresh weight of 10-day-old seedlings of the indicated genotypes treated with indicated UVC doses as in (C). Fresh weight was normalized to the untreated (0 J/m^2^) samples of the same genotype. n = 3 independent replicates. (E) Log2 fold change of gene expression of UVC-inducible CAMTA3 target genes in indicated genotypes at indicated time points. (F) Heatmap showing the fold enrichment of the top 15 GO terms enriched in UVC-inducible CAMTA3 target genes. For (D) and (E), data show means ± SD. Different letters mean *p*<0.05 for ANOVA analysis followed by Fisher’s LSD posthoc test.

We next asked whether CRYs and UBP12/13 would regulate the DDR through CAMTAs as well. To test this hypothesis, we generated DREM models using expression of the DEGs derived from *cry1cry2* and *ubp12ubp13* (Figures S3C and S3D) along the time course. In both the *cry1cry2* and *ubp12ubp13* models, SOG1 and WRKYs were predicted to regulate gene expression after UVC (Figures S3C and S3D). However, CAMTAs were only predicted in the *ubp12ubp13* but not in the *cry1cry2* DREM model (Figures S3C and S3D). This suggests that CAMTAs may be dysregulated in the *cry1cry2* mutant. To explore this possibility, we compared the expression of CAMTA3 target genes in *cry1cry2* and *ubp12ubp13* to WT, respectively. We obtained UVC-inducible CAMTA3 target genes by overlapping the upregulated genes in WT after UVC in our RNA-seq data with targets of CAMTA3 previously identified by chromatin immunoprecipitation sequencing (ChIP-seq) (Table S2)^57^. Compared to WT, these CAMTA3 target genes were less induced in *cry1cry2* and more induced in *ubp12ubp13* at 15 min and 60 min after UVC (Figure 3E, Table S2). Interestingly, the UVC-inducible CAMTA3 target genes are enriched in GO terms related to stress responses (Figure 3F, Table S3), which is similar to the upregulated GO terms in WT after UVC (Figure S2D), suggesting that CRYs and UBP12/13 may antagonistically regulate the DNA damage-induced transcriptional changes partially through the CAMTAs.

### UVC induces the UBP12/13-dependent CRY2 degradation

We found that UBP12/13 and CRYs had opposite functions in the response to UVC (Figures 1B, 1E and 2C), which is reminiscent of the opposite roles of UBP13 and CRY2 in blue light signaling pathway^25^, where UBP13 interacts with CRY2 in a blue light-dependent manner^25^. Therefore, we asked whether UVC could also enhance the interaction between CRY2 and UBP13, similar to blue light. To address this, we treated 5-day-old seedlings expressing both FLAG-CRY2 and UBP13-HA with or without UVC and performed co-immunoprecipitation (co-IP) analysis using a FLAG antibody. The UVC light was obtained through a light-emitting diode (LED) lamp, which did not contain UVA or blue light (Figure S4A). This UVC LED lamp also induced the hypersensitive phenotype of the *cry1cry2* mutant (Figure S4B). We found that compared to untreated samples, UVC-treated seedlings exhibited enhanced interaction between CRY2 and UBP13 (Figure 4A). This result suggests that UVC can strengthen the interaction between CRY2 and UBP13.

**Figure 4.**
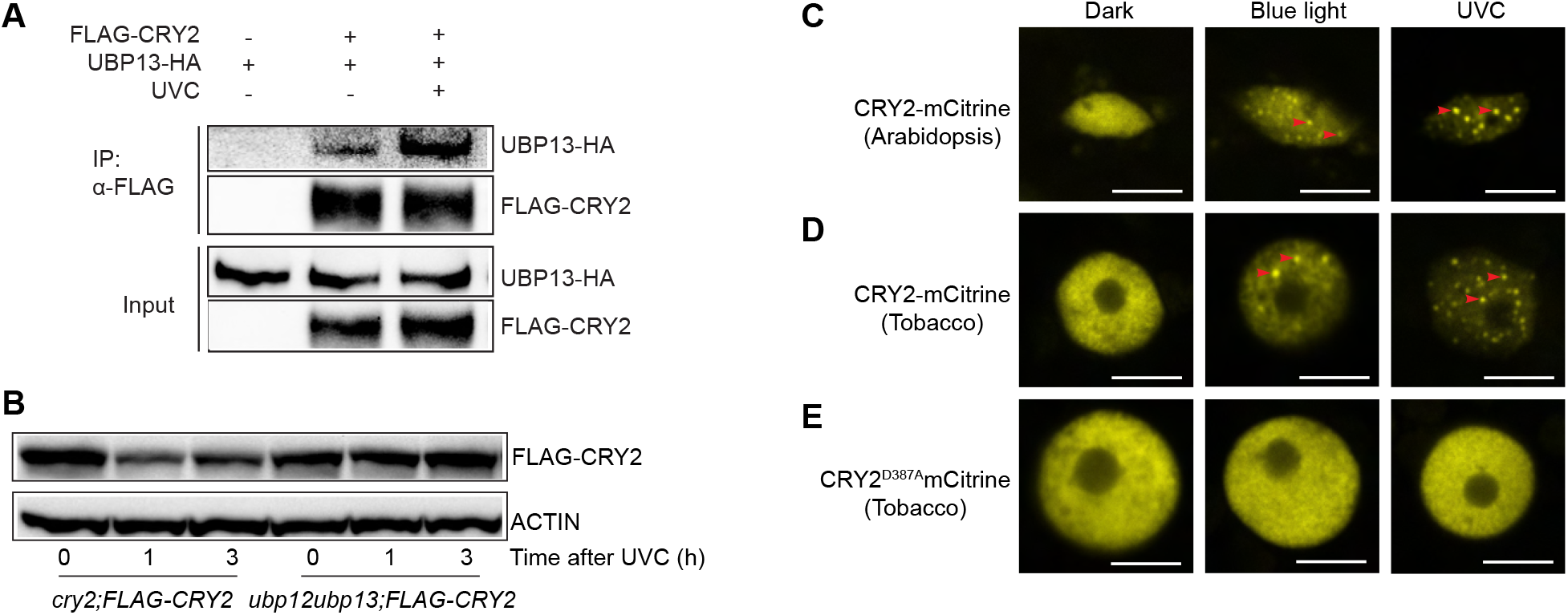
CRY2 interaction with UBP13 and nuclear speckle formation is induced by UVC. (A) Co-IP immunoblot showing enhanced pulldown of UBP13-HA by FLAG-CRY2 after UVC treatment. 4-day-old light-grown seedlings expressing FLAG-CRY2 and UBP13-HA were dark adapted for 24 h, then treated or untreated with approximately 1800 J/m^2^ of UVC from the LED source and collected after 10 min incubation in the dark. Seedlings expressing only UBP13-HA without UVC treatment were used as a negative control for the co-IP. (B) Immunoblot analysis of FLAG-CRY2 levels after UVC treatment in the *cry2* and *ubp12ubp13* mutant backgrounds. 5-day-old light-grown seedlings were treated with 6000 J/m^2^ of UVC and collected after 0, 1, and 3 h. (C-E) Representative confocal microscopy images of nuclei in plants expressing CRY2-mCitrine (C, D) or CRY2^D387A^-mCitrine (E) fusions in the dark and after blue light or UVC in transgenic *Arabidopsis* seedlings (C) and infiltrated *Nicotiana benthamiana* leaves (D, E). Samples were fixed before imaging. The scale bar is 5 μm. For (C-D), red arrowheads indicate representative nuclear speckles.

Blue light-enhanced interaction between UBP13 and CRY2 promotes COP1-mediated CRY2 degradation^25^, therefore, we next asked whether UVC also induces CRY2 degradation. To test this hypothesis, we treated 5-day-old *cry2* mutant seedlings expressing FLAG-CRY2 with UVC and analyzed FLAG-CRY2 protein levels after 0, 1, and 3 h. We found that FLAG-CRY2 protein levels diminished 1 h after UVC and partially recovered 3 h after UVC (Figures 4B and S4C), suggesting that UVC can induce CRY2 degradation. Next, we asked whether UVC-induced CRY2 degradation was dependent on UBP12/13. We treated *ubp12ubp13* seedlings expressing FLAG-CRY2 with UVC, and found that FLAG-CRY2 was more stable in the *ubp12ub13* mutant background than in the *cry2* mutant background in UVC (Figures 4B and S4C). This result suggests that UVC-induced CRY2 degradation is partially dependent on UBP12/13. In blue light, UBP12/13 regulates CRY2 degradation through COP1^25^. Therefore, we next examined whether COP1 plays a role in UVC response. To address this hypothesis, we treated *cop1-4* and *COP1oe* seedlings with UVC and found that *cop1-4* is hyposensitive while *COP1oe* is hypersensitive to DNA damage (Figures S4D and S4E), suggesting that COP1, similar to UBP12/13, promotes plant resistance against UVC-induced DNA damage. Together, these results indicate that UBP12/13 may regulate DDR by interacting with CRY2 and modulating CRY2 degradation.

### UVC induces the formation of CRY2 nuclear speckles

Upon exposure to blue light, CRY1 and CRY2 form punctate nuclear speckles, also known as photobodies^18,19^. Since we found that UVC could enhance the CRY2-UBP13 interaction and induce UBP12/13-dependent CRY2 degradation, similar to blue light, we next examined if UVC could also induce CRY2 speckles. We treated 4-day-old dark-grown *cry2;CRY2-mCitrine* and *cry2;mCitrine-CRY2* seedlings with blue light, UVC light from the LED lamp or continued darkness to observe speckle formation of CRY2. To excite mCitrine in confocal microscopy, a blue light source is often used^58^. Therefore, to prevent microscopy-induced CRY2 speckles, we used a mild fixative to crosslink proteins in seedlings before confocal imaging^18^. C-terminally tagged CRY2, such as CRY2-GFP, is known to readily form speckles in blue light while N-terminally tagged CRY2, like GFP-CRY2, requires blocking of proteasome-mediated protein degradation to form speckles in blue light^18^. Therefore, we first examined the formation of speckles of CRY2-mCitrine in *Arabidopsis* seedlings. As expected, in darkness we did not observe CRY2-mCitrine speckles (Figures 4C and S4F), while in blue light (40 μmol m^-2^ s^-1^) and UVC (approximately 1800 J/m^2^) speckles formed after 2 min (Figures 4C and S4F). Next, we examined whether mCitrine-CRY2 could also form speckles under UVC. mCitrine-CRY2 did not form speckles in the dark (Figure S4G), in contrast, it formed nuclear speckles after 30 min of blue light (40 μmol m^-2^ s^-1^) and under UVC (30 min; approximately 100 J/m^2^) (Figure S4G). Together, these results suggest that UVC can induce the formation of CRY2 nuclear speckles.

The ability of CRY2 to absorb blue light to form speckles is dependent on its covalently bound chromophore, flavin adenine dinucleotide (FAD)^59,60^. To test if CRY2 requires its light-sensing property to form speckles under UVC, we generated CRY2^D387A^ where aspartic acid 387 was substituted with alanine within the FAD-binding pocket, rendering it light-insensitive^61^. mCitrine-CRY2^D387A^ did not form speckles in dark, blue light, or UVC (Figure S4H), suggesting that CRY2 requires its light-sensing ability to form nuclear speckles under UVC. To check whether CRY2 forms nuclear speckles in UVC when expressed in a heterologous system, we transiently expressed CRY2-mCitrine as well as CRY2^D387A^-mCitrine in *Nicotiana benthamiana*. We found that CRY2-mCitrine formed nuclear speckles under blue light and UVC (Figure 4D), while CRY2^D387A^-mCitrine remained uniformly distributed in the nucleus in dark, blue light and UVC (Figure 4E). This result strengthens the conclusion that *Arabidopsis* CRY2 requires its light-sensing activity to form nuclear speckles under UVC. Taken together, our data suggested that UVC induces similar changes in CRY2 as blue light, including enhancing the interaction between CRY2 and UBP13, inducing CRY2 degradation and triggering CRY2 nuclear speckle formation.

## DISCUSSION

In this study, we demonstrate that DDR is favorably regulated by CRYs in *Arabidopsis.* We also show that UBP12 and UBP13 negatively regulate many aspects of CRY-mediated DDR, including CPD repair and induction of DNA repair genes (Figures 1 and S1). Through transcriptomic analysis, we found that CRYs and UBP12/13 antagonistically regulate the transcriptional response to DNA damage (Figures 2 and S2) and identified CAMTA transcription factors as novel regulators of DDR (Figures 3 and S3). Upon further investigation, we unexpectedly discovered that CRY2 responds to UVC in a manner similar to blue light, such as interacting stronger with UBP13, undergoing UBP12/13-dependent degradation, and forming nuclear speckles (Figures 4 and S4). Together, our results reveal key roles for CRYs and UBP12/13 in the DDR and suggest a mechanism where UBP12/13 destabilizes CRY2 during the DDR.

Evolved from photolyases, present-day CRYs have lost their enzymatic activity to repair pyrimidine dimers^30^, however, they still bind to damaged DNA^31^, indicating that although CRYs cannot directly repair UV-damaged DNA, they might have a residual function in sensing or responding to DNA damage^32^. There is evidence in mammals in favor of this hypothesis, as DNA damage affects CRY protein stability: CRY1 is stabilized, while CRY2 is destabilized^32,62.^. The roles of CRYs in DDR can also differ between paralogs *(i.e.* CRY1 and CRY2) and homologs *(e.g.*, human and mouse)^32,62^. For instance, in human cell lines, stabilized CRY1 promotes DNA repair by regulating genes involved in HR repair of DSBs^62^, while mouse CRY1 can function as a transcriptional repressor^32^. Mouse CRY2 inhibits the transcription of DNA damage responsive genes, therefore, destabilization of CRY2 upon DNA damage releases gene expression and induces DNA damage response^32^. Here we show that upon UVC-induced DNA damage, plant CRY2 proteins are destabilized, as in mice^62^, and that that CRY1 and CRY2 together promote DNA repair by regulating the transcription of genes involved in HR, as well as RAD51 protein (Figures 1G, S1G-H), which is similar to the role of human CRY1^62^. Therefore, our study suggests that not only animal CRYs, but also plant CRYs play a residual role in DDR. Further investigations would be required to address the effects of UVC on CRY1 stabilization in plants and the its functional consequences to DDR.

In mammalian cells, CPDs are mainly repaired by the NER pathway, which removes one strand of DNA containing the damaged site and replaces it with newly synthesized DNA^12^. Unlike mammals, plants have the PHR1 photolyase, which uses energy from light to efficiently repair CPD without DNA excision^63^. For this reason, it was thought that photolyase-dependent repair in plants was the major repair pathway of CPDs in light conditions and the NER repair was only relevant in the dark^64^. However, a recent study suggests that both photolyases and the NER pathway are important for repairing UV-induced DNA damage in light, as there is a synergistic genetic interaction between *PHR1* and the NER-related *CUL4, DDB1A* and *DDB2*^64^. Our study finds that CRYs promote the repair of CPDs under light (Figures 1C-E, S1E). In this context, CRYs may regulate the repair of CPDs either by PHR1 photolyase-mediated repair or by NER. On one hand, CRYs may regulate the expression of *PHR1* through the light signaling pathway. For instance, the TF ELONGATED HYPOCOTYL 5 (HY5) induces the expression of *PHR1* in light^40,65^, but is in turn repressed by another light signaling component, DE-ETIOLATED 1 (DET1)^40^. Thus, since CRYs are known to positively regulate HY5 protein stability^66^, it is plausible that CRYs indirectly induce *PHR1* via HY5 under UVC, which is consistent with our observation that CRYs promote CPD repair, opposite to DET1^40^. On the other hand, it is also plausible that CRYs regulate NER through the CRL4^COP1/SPA^ complex. First, CRYs can repress the activity of COP1^66,67^, which we found is a negative regulator of the DDR (Figures S4D-E). Second, similar to COP1, DET1 forms a complex with CUL4 and DDB1 to regulate NER in collaboration with DDB2^68^. Therefore, the CRL4^COP1/SPA^ complex may serve as a mediator between CRY-mediated light signaling and the NER-mediated repair of CPDs. In addition, the finding that CRYs are not required for DSB repair (Figure S1D) further suggests that photolyase-mediated repair and the NER are the two most plausible DNA repair pathways that could be regulated by CRYs.

DDR, however, isn’t regulated just at the transcriptional level. Proteins involved in DDR are regulated by post-translational modifications (PTMs)^69^. After phosphorylation, ubiquitination is the second most prevalent PTM^70^, which alters protein stability and protein-protein interactions. For example, p53 is destabilized by ubiquitination^71^. Moreover, ubiquitination of histone H2AX promotes the recruitment of DNA repair proteins to DNA damage sites^72^. DUBs, proteases that remove ubiquitination from target proteins, play an important role in animal DDR, for instance, USP7, the ortholog of UBP12/13 in animals, stabilizes p53^73^ and the Chk1 kinase^74^, which is essential for the initiation of the DDR^75^. However, there are only a few papers exploring the role of DUBs in plant DDR. Recently, Al Khateeb et al. suggest that UBP12, a plant DUB, acts as a positive regulator of UVC tolerance in the dark^76^. In contrast to their study, our study finds that UBP12/13 act as negative regulators of DDR in light conditions (Figures 1, 2, S1 and S2). This variation in results may arise from the difference in experimental procedures, suggesting that the function of UBP12/13 is distinct in the light versus dark. Although our study suggests that UBP12/13 likely regulate the DDR through CRYs (Figure 4A-B), we cannot rule out the possibility that UBP12/13 could target other DDR-related proteins. For instance, UBP12/13 can deubiquitinate histone H2A^77^. Because histone ubiquitination marks are important signals in DDR for the recruitment of DNA damage repair proteins^78^, it is plausible that the negative role of UBP12/13 in plant DDR could also result from removing of histone ubiquitination marks.

DDR promotes DNA damage repair, inhibits the cell cycle to allow sufficient time for DNA repair, and induces apoptosis in cells that have irreparable DNA damage^79^. Constitutive activation of the latter two aspects of DDR in the absence of DNA damage could lead to undesired cell cycle arrest and cell death^71^. Therefore, organisms evolved mechanisms to desensitize the DDR. For example, p53 can induce the expression of its E3 ligase, Mdm2, which in turn leads to p53 degradation, serving as a negative feedback loop to halt the p53 signaling pathway when DNA damage is repaired^71^. This inhibition of p53 by Mdm2 is also important for normal cell survival as Mdm2-deficient mice are embryonically lethal due to the cytotoxicity caused by ectopic activation of p53^80^. Similarly, we show that UBP12/13 serve as a brake for CRY-mediated DDR in plants (Figures 1-2, S1-2, 4A-B). This inhibition of CRY function by UBP12/13 is also crucial for normal plant growth since the loss of UBP12 and UBP13 leads to over-accumulation of CRY2 and subsequently constitutive activation of stress responses resulting in stunted growth phenotypes^25^.

CAMTA transcription factors are conserved across many animal and plant species^81^. In animals, CAMTAs regulate nervous system-related processes and cardiac growth^82^–^85^, while in plants, they are mainly implicated in abiotic stress and immune responses^56^. In both groups, the role of CAMTAs in DDR remains unexplored. Our study shows that CAMTA TFs play a novel role in UVC-induced DDR (Figures 3, S3). Many TFs are important for DDR, especially SOG1, as gamma irradiation-induced gene expression is largely diminished in the *sog1* loss of function mutant ^11,86^. However, using public databases^87^, we found that induction of the *CAMTA3* gene 20 min after gamma irradiation is largely unaffected by the genetic loss of *SOG1*^11^, suggesting that CAMTAs might play a SOG1-independent role in the DDR, similar to E2Fa, a known SOG1-independent TF^88^. CAMTAs can bind to calmodulin proteins that are important for the calcium signaling pathway^56^. In animals, the calcium signaling pathway is required for the DDR. Intracellular calcium level is increased upon DNA replication stress, which in turn activates the calcium signaling pathway and inhibits Exonuclease 1 (Exo1) from making aberrant nicks in replication forks, thus maintaining genome stability^89^. Therefore, further investigations would provide insights into whether calcium signaling play a role in plant DDR and if this role is dependent on the CAMTAs and CRYs.

CRYs are well-characterized blue/UVA light receptors^90,91^, and evidence suggests that human CRY1 and the chromophore common to all CRYs, FAD, can absorb light in the UVB/UVC spectrum^92,93^. However, whether UVC light is functionally relevant for CRYs has never been explored. Blue light triggers the formation of CRY2 nuclear speckles^18^, where photoactivated CRY2 carries out its function^20^. Our unexpected finding that CRY2 requires its light-sensing property to form nuclear speckles in UVC (Figures 4 and S4) suggests that this light stimulus could trigger the photoactivation of CRY2. This discovery not only provides the first evidence that the UVC light spectrum is functionally relevant for CRYs, but also justifies future research to explore if CRY2 could act as a bona fide UVC light receptor, with experiments to be performed such as the spectroscopic examination of CRY2 upon UVC exposure.

Recent studies have shown that CRYs and the UVB receptor, UVR8, functionally interact^33–35^. Even though *cry1, cry1cry2* and *uvr8* mutants survive under natural and simulated sunlight (*i.e.*, supplemented with UVB), *cry1uvr8* double and *cry1cry2uvr8* triple mutants do not, suggesting that CRYs and UVR8 redundantly contribute to plant survival in sunlight^33–35^. Paradoxically, CRY proteins induce the dimerization and, therefore, inactivation of UVR8 in a blue-light-dependent manner^33–35^. Moreover, CRYs likely oppose UVR8-induced gene expression under UVB, suggesting that CRYs would function as a brake to UVR8 hyper-activation^33–35^. In this context, the mechanism of how CRYs positively contribute to plant survival under UVB remains largely unknown. In our study, we further extended the function of CRYs into the UVC spectrum and showed that CRYs play a role in UVC-induced DDR, and presented evidence that CRYs could regulate DNA repair after UVC exposure to contribute to plant growth and survival. Therefore, beyond revealing a novel role for CRYs, UBP12/13 and CAMTAs in UVC, our findings might point to how CRYs help plants to survive under other types of genotoxic stresses, such as UVB.

## ACKNOWLEDGEMENTS

We thank Dr. Julie Law for the *ku70* seeds and Dr. Michael Thomashow for the *camta123* seeds. We thank Dr Laura Taylor for critical reading of this manuscript. This work is supported by National Institutes of Health (NIH) grant R35GM125003 to U.V.P. NIH GM12500303S1 and GM12500304S1 grant for the purchase of confocal microscope is acknowledged by U.V.P. The George A. and Marjorie H. Anderson fellowship is acknowledged by Y.H.

## AUTHOR CONTRIBUTIONS

U.V.P. conceived the study. Y.H., L.N.L. and U.V.P. designed all the experiments and Y.H. performed most of the experiments except the following: Y.H. and L.N.L performed the UVC fresh weight experiments. L.N.L performed the qPCR experiments to examine DNA repair marker gene expression. Y.H. and D.R. performed RNA-seq analysis, Y.H., L.N.L, and J.M. performed molecular cloning and generated transgenic lines, Y.H., D.R. and U.V.P. wrote the manuscript, and all authors reviewed and revised the manuscript.

## MATERIALS AND METHODS

### Plant genotypes and growth conditions used

*Arabidopsis thaliana Columbia* (*Col-0*) ecotype was used as background for mutants and transgenic lines. *cry1-304*^94^, *cry2-1*^91^, *cry1-304 cry2-1*^94^, *ubp12-2w ubp13-3*^36^, *ku70*^95^, *cop1-4*^96^ and *camta123*^97^ mutants have been previously described. *Col-0;UBQ10_pro_:UBP13-6xHA* (*UBP13oe*)^25^, *cry2-1;UBQ 10_pro_:UBP13-6xHA;CRY2_pro_:2xStrep-6xHis-3xFLAG-CRY2*^25^, *ubp12-2w ubp13-3;UBQ10_pro_: 2xStrep-6xHis-3xFLAG-CRY2 (ubp12ubp13;CRY2oe)*^25^, *Col-0;UBQ10_pro_:COP1-6xHis-3xFLAG (COP1oe)*^25^, and *cry2-1;UBQ10_pro_:2xStrep-6xHis-3xFLAG-CRY2* (*cry2;CRY2oe*)^98^ lines have been described previously. After surface sterilization, seeds were plated on 0.5× Linsmaier and Skoog (LS) medium (HiMedia Laboratories) containing 0.8% agar, stratified for 2 days in darkness at 4°C and then grown at 22°C under 100 μmol m^-2^ s^-1^ white light from a LED source in a growth chamber (Percival Scientific) unless otherwise specified.

### Molecular cloning and transformation of *Arabidopsis* and *Nicotiana benthamiana*

Promoters and coding sequences were amplified from genomic or cDNA pool of Col-0 (WT) plants or subcloned from plasmids by PCR and cloned into Gateway donor plasmids including pDONR221, pDONRP4-P1R and pDONRP2R-P3 (Thermo Fisher Scientific) using BP Clonase II (Thermo Fisher Scientific). CRY2^D387A^ was generated by replacing the aspartate 387 with an alanine through site-directed mutagenesis (Oligos listed in Table S4). Three-fragment Gateway cloning technology was used to combine the Gateway donor constructs with pB7m34GW or pK7m34GW destination plasmids^99^ using LR Clonase II (Thermo Fisher Scientific). Binary destination plasmids were transformed into *Agrobacterium tumefaciens* (GV3101) and then transformed into *Arabidopsis* plants using the floral dip method^100^. *cry1-304 cry2-1;UBQ10_pro_:UBP13-6xHA (cry1cry2;UBP13oe)* line was generated by transforming *cry1cry2* with *pB7m34GW-UBQ10_pro_:UBP13-6xHA* plasmid. *cry2* plants were transformed either with *pK7m34GW-UBQ10_pro_:CRY2-mCitrine, pB7m34GW-CRY2_pro_:mCitrine-CRY2* or *pB7m34GW-CRY2_pro_:mCitrine-CRY2^D387A^* plasmids to generate *cry2;CRY2-mCitrine, cry2;mCitrine-CRY2* or *cry2;mCitrine-CRY2D^387A^*, respectively. *pK7m34GW-UBQ10_pro_:CRY2-mCitrine* and *pB7m34GW-UBQ10_pro_:CRY2^D387A^-mCitrine* were transformed into *Agrobacterium* and used to infiltrate *Nicotiana benthamiana* plants as described before^25^.

### UVC sensitivity assay

UVC treatment was performed using a UV Crosslinker 1800 (Stratagene) or with a UVC-emitting LED lamp (peak wavelength 270-280 nm), (Cat# E275-80-Module; International Light Technologies). Plants were grown in continuous white light for 4 days at 22°C, then treated with 5500 or 8000 J/m^2^, and returned to continuous white light for another 5-6 days before phenotyping and measurement of fresh weight. Three biological replicates of fresh weight measurement were performed. For each biological replicate, the total fresh weight of 10-24 seedlings was measured and the fresh weight per seedling was calculated. Fresh weight percentages were calculated by normalizing the fresh weight measurement at the indicated UVC dose to the 0 J/m^2^ treatment group of the same genotype.

### Zeocin sensitivity assay

Plants were grown under long days (LD) for 4 days, then transferred to plates containing 0, 4 or 8 μM of zeocin (Thermo Fisher Scientific) and grown for further 8 days in LD before fresh weight measurement. Three biological replicates of fresh weight measurements were performed. Fresh weight and fresh weight percentage relative to 0 μM were calculated as described above.

### CPD dot blot assay

5-day-old seedlings grown in LD were treated with 6000 J/m^2^ UVC using the UV crosslinker and then transferred to 100 μmol m^-2^ s^-1^ white light for 1 min or 180 min before flash freezing in liquid nitrogen. Genomic DNA was extracted with the cetyltrimethylammonium bromide (CTAB) method^101^, denatured by incubating at 100°C for 10 min and placed on ice immediately for 15 min, and quantified using a Qubit fluorometer (Thermo Fisher Scientific) and a Qubit ssDNA assay kit (Thermo Fisher Scientific). Serial dilutions (1, 1:10, 1:100) of the genomic DNA were blotted onto a Whatman Nytran SuPerCharge nylon blotting membrane (MilliporeSigma) and baked at 80°C for 2 h, then soaked in tris-buffered saline (TBS) with Tween-20 (TBST) (20 mM Tris-HCl, 150 mM NaCl, 0.05% Tween-20, pH 7.6) for 20 min before blocking with 5% fat-free milk in TBST for 30 min. After blocking, the membrane was incubated with an anti-CPD antibody (Cosmo Bio USA) at 1:1000 dilution prepared in 1% fat-free milk made in TBST overnight at 4°C before washing with TBST three times, 5 min each. Following the wash, the membrane was incubated with 1:10,000 dilution of anti-mouse-horseradish peroxidase (anti-mouse-HRP) antibody (Bio-Rad) at room temperature for 1 h and washed again three times in TBST, 5 min each wash. Imaging was performed in a Chemidoc imaging system (Bio-Rad) following the addition of SuperSignal West Dura Extended Duration Substrate (Thermo Fisher Scientific) to the membrane. Methylene blue staining was performed by incubating the blotted and baked nylon membrane with staining buffer (0.04% methylene blue, 0.5 M sodium acetate, pH 5.2) for 10 min, then de-stained with distilled water for 5 min before imaging. The CPD dot blots were quantified with ImageJ^102^.

### Protein extraction and Immunoblotting

Total protein was extracted by grinding frozen *Arabidopsis* tissue in lithium dodecyl sulfate (LDS) buffer (106 mM Tris-HCl, 141 mM Tris, 2% LDS, 10% glycerol, 0.51 mM EDTA, 0.22 mM Coomassie Brilliant Blue G 250 (SERVA Electrophoresis GmbH), 0.175 mM phenol red, 10% tris(2-carboxyethyl)phosphine). After centrifugation at 13000 rpm for 5 min, proteins were separated by sodium dodecyl-sulfate polyacrylamide gel electrophoresis (SDS-PAGE) in either homemade 7% or 4-12% gradient Bis-Tris polyacrylamide gel (Thermo Fisher Scientific) using 3-(N-morpholino)propanesulfonic acid (MOPS) running buffer (40 mM MOPS, 10 mM sodium acetate, 1 mM EDTA, pH 7) and transferred to nitrocellulose membrane (MilliporeSigma). After transfer, the nitrocellulose membrane was incubated in 5% fat-free milk made in TBST for 30 min, followed by incubation with primary antibodies in 1% fat-free milk made in TBST for 1 h. Then the membrane was washed three times with TBST and incubated with the secondary antibodies in 1% fat-free milk made in TBST for 1 h. The blots were washed three times with TBST and detection was performed as described above. The following antibodies were used: anti-RAD51 (Cat# AB63799, Abcam), anti-phospho-p44/42 MAPK (Erk1/2) (Thr202/Tyr204) (Cell Signaling Technology), anti-actin (MP Biomedicals) as primary antibodies. Goat-anti-mouse-HRP (Bio-Rad) and goat-anti-rabbit-HRP (Bio-Rad) were used as secondary antibodies. Conjugated anti-HA-HRP (Cat#12013819001, MilliporeSigma) and anti-FLAG-HRP (Thermal Fisher Scientific) antibodies were used to detect HA- and FLAG-tagged proteins, respectively. All immunoblot experiments were repeated at least twice. Quantification of the immunoblot was performed in ImageJ software^102^ by measuring the mean gray value of bands subtracted by the mean gray value of the background.

### *In vivo* co-immunoprecipitation

4-day-old *cry2-1;UBQ10_pro_:UBP13-6xHA;CRY2_pro_: 2xStrep-6xHis-3xFLAG*-CRY2 seedlings grown under continuous white light were dark adapted for 24 h, then treated with continued darkness or UVC LED source (approximately 1800 J/m^2^). Tissue was collected after 10 min, immediately frozen and later ground in liquid nitrogen. Each 1 g of tissue was dissolved in 2 ml of SII buffer (100 mM sodium phosphate [pH 8], 150 mM NaCl, 5 mM EDTA, 5 mM EGTA, 0.1% Triton X-100, 1× protease inhibitors (Sigma), 50 μM MG132) and sonicated (Branson Ultrasonics) on the ice at 40% power, with 0.5 s on/off cycles for a total of 10 s. The protein extracts were then clarified by two rounds of centrifugation at 13000 rpm for 10 min at 4°C. Protein concentration was inferred by spectroscopy using Bradford reagent (Bio-Rad), and normalized for inputs and co-IPs. For co-IPs, proteins were then mixed with anti-FLAG antibody (Thermal Fisher Scientific) for 1 h at 4 °C and incubated with protein-G magnetic beads (Bio-rad) for 0.5 h at 4 °C. Beads were washed 3× with 0.75 ml of SII buffer and proteins were eluted with 20 μl of 2× LDS buffer and boiled at 95°C for 5 min before immunoblot analysis, as described above.

### Laser scanning confocal microscopy

For CRY2-mCitrine, 4-day-old dark-grown *Arabidopsis cry2;UBQ10_pro_::CRY2-mCitrine* seedlings were incubated in MG132 buffer (0.5× LS medium, 50 μM MG132) for 5-8 h in the dark at room temperature. Samples were then treated with 40 μmol m^-2^ s^-1^ of blue light or UVC LED (approximately 1800 J/m^2^ in total) for 2 min or kept in continued darkness. Seedlings were immediately fixed in 4% PFA with a vacuum for 20 min, then washed for 5 min twice in phosphate-buffered saline (PBS) before cotyledon cells were imaged. For mCitrine-CRY2 and mCitrine-CRY2^D387A^, 4-day-old dark-grown *cry2;CRY2_pro_::mCitrine-CRY2* or *cry2;CRY2_pro_::mCitrine-CRY2^D387A^* seedlings were incubated in MG132 buffer (0.5× LS medium, 50 μM MG132) for 0.5 h in the dark at room temperature. Then treated for 30 min with 40 μmol m^-2^ s^-1^ of blue light or UVC LED (approximately 100 J/m^2^ in total) or continued darkness. Seedlings were fixed in 1% PFA for 10 min and washed twice in PBS for 5 min. Hypocotyl cells were imaged in this case.

*UBQ10_pro_::CRY2-mCitrine* or *UBQ10_pro_::CRY2^D387A^-mCitrine* were transiently expressed in *N. benthamiana* following agroinfiltration of leaves. For this, plants were grown for approximately four weeks in the greenhouse environment. Three leaves were infiltrated per condition, and then plants were kept in white light for 1 day and dark incubated for 2 days to allow for CRY2 accumulation. After this time, leaves were infiltrated with 50 μM MG132 for 0.5 h, prior to treatment with blue light (40 μmol m^-2^ s^-1^) or UVC LED (approximately 4500 J/m^2^ in total) for 5 min or continued darkness. Leaves were fixed in 4% PFA for 20 min with a vacuum, then kept in 1× PBS and imaged under the confocal microscope. All confocal microscopy was performed using the LSM900 confocal microscope (Zeiss) using a 488 nm laser and images were captured at the emission range of 410 to 545 nm.

### mRNA sequencing and analysis

5-day-old seedlings grown under LD conditions were untreated (0 min) or treated with 6000J/m^2^ UVC and collected after 15, 60 and 180 min. Two biological replicates were harvested for each sample and frozen in liquid nitrogen. RNA was extracted using a Direct-zol RNA miniprep kit (Zymo Research) and quantified using a Qubit fluorometer (Thermo Fisher Scientific). 500 ng of total RNA was used for mRNA isolation using NEBNext poly(A) mRNA Magnetic Isolation Module (New England Biolabs) and the purified mRNA was used to construct libraries using the NEBNext Ultra II Directional RNA Library Prep Kit for Illumina (New England Biolabs) following manufacturer instructions. Single-end sequencing of 76 bp was performed on NextSeq500 (Illumina) to a total of 40 million reads per sample on average. The sequencing reads were mapped to the *Arabidopsis thaliana* Col-0 genome (TAIR10) using STAR version 2.7.5c^103^. Differential gene expression analysis was performed using Cufflinks version 2.2.1^104^. R environment version 4.1.0 (R Foundation) and its packages (ggplot2, RColorBrewer, corrplot, DESeq2^105^) were used for statistical analysis and to visualize the results. Principal component analysis was performed using DESeq2^105^. GO term analysis was performed using PANTHER^106^.

### RT-qPCR analysis

Total RNA was extracted from frozen *Arabidopsis* seedlings using the Direct-zol RNA miniprep kit (Zymo Research). cDNA from RNA was synthesized using the iScript cDNA Synthesis Kit (Bio-Rad) and qPCR was performed using the indicated oligos (Table S4) (QuantStudio 6 Pro PCR system; Thermo Fisher Scientific) using Power SYBR Green Master Mix (Thermo Fisher Scientific). Expression values were normalized to the *UBC28* reference gene and calculated using the 2^-ΔΔCt^ method^107^.

### DREM analysis

DREM analysis was performed as previously described^11^. For each genotype (WT, *cry1cry2* and *ubp12ubp13*), the log2 fold change of the expression of all the DEGs along the time course in the corresponding genotype was used as inputs for DREM models^53^. The TF-gene interaction file derived from Bourbousse et al^11^ was used as input for the DREM analysis.

### Discovery of *de novo* motifs

The *de novo* motif search by HOMER^54^ was performed using lists of target genes (genes within the W2 group from the DREM model, all upregulated genes in WT after UVC, and all downregulated genes in WT after UVC) as input. The following code was used in a Linux environment: “findMotifs.pl /file/path/to/gene/names arabidopsis /file/path/to/output -noconvert -start -500 -end 50 - nogo”.

## REFERENCES

1. Manova, V., and Gruszka, D. (2015). DNA damage and repair in plants – From models to crops. Front Plant Sci 6, 1–26. 10.3389/fpls.2015.00885.

2. Alhmoud, J.F., Woolley, J.F., al Moustafa, A.-E., and Malki, M.I. (2020). DNA Damage/Repair Management in Cancers. Cancers (Basel) 12, 1050. 10.3390/cancers12041050.

3. Roldán-Arjona, T., and Ariza, R.R. (2009). Repair and tolerance of oxidative DNA damage in plants. Mutation Research/Reviews in Mutation Research 681, 169–179. 10.1016/j.mrrev.2008.07.003.

4. Gill, S.S., Anjum, N.A., Gill, R., Jha, M., and Tuteja, N. (2015). DNA Damage and Repair in Plants under Ultraviolet and Ionizing Radiations. The Scientific World Journal 2015, 1–11. 10.1155/2015/250158.

5. Pfeifer, G.P., You, Y.H., and Besaratinia, A. (2005). Mutations induced by ultraviolet light. Mutation Research - Fundamental and Molecular Mechanisms of Mutagenesis 571, 19–31. 10.1016/j.mrfmmm.2004.06.057.

6. Groelly, F.J., Fawkes, M., Dagg, R.A., Blackford, A.N., and Tarsounas, M. (2022). Targeting DNA damage response pathways in cancer. Nat Rev Cancer. 10.1038/s41568-022-00535-5.

7. Savitsky, K., Bar-Shira, A., Gilad, S., Rotman, G., Ziv, Y., Vanagaite, L., Tagle, D.A., Smith, S., Uziel, T., Sfez, S., et al. (1995). A Single Ataxia Telangiectasia Gene with a Product Similar to PI-3 Kinase. Science (1979) 268, 1749–1753. 10.1126/science.7792600.

8. Weinert, T.A., Kiser, G.L., and Hartwell, L.H. (1994). Mitotic checkpoint genes in budding yeast and the dependence of mitosis on DNA replication and repair. Genes Dev 8, 652–665. 10.1101/gad.8.6.652.

9. Smith, J., Mun Tho, L., Xu, N., and A. Gillespie, D. (2010). The ATM–Chk2 and ATR–Chk1 Pathways in DNA Damage Signaling and Cancer. In, pp. 73–112. 10.1016/B978-0-12-380888-2.00003-0.

10. Linzer, D.I.H., and Levine, A.J. (1979). Characterization of a 54K Dalton cellular SV40 tumor antigen present in SV40-transformed cells and uninfected embryonal carcinoma cells. Cell 17, 43–52. 10.1016/0092-8674(79)90293-9.

11. Bourbousse, C., Vegesna, N., and Law, J.A. (2018). SOG1 activator and MYB3R repressors regulate a complex DNA damage network in Arabidopsis. Proc Natl Acad Sci U S A 115, E12453–E12462. 10.1073/pnas.1810582115.

12. Marteijn, J.A., Lans, H., Vermeulen, W., and Hoeijmakers, J.H.J. (2014). Understanding nucleotide excision repair and its roles in cancer and ageing. Nat Rev Mol Cell Biol 15, 465–481. 10.1038/nrm3822.

13. Lee, J., and Zhou, P. (2007). DCAFs, the Missing Link of the CUL4-DDB1 Ubiquitin Ligase. Mol Cell 26, 775–780. 10.1016/j.molcel.2007.06.001.

14. Sancar, A. (1994). Structure and function of DNA photolyase. Biochemistry 33, 2–9. 10.1021/bi00167a001.

15. Chaves, I., Pokorny, R., Byrdin, M., Hoang, N., Ritz, T., Brettel, K., Essen, L.O., van der Horst, G.T.J., Batschauer, A., and Ahmad, M. (2011). The cryptochromes: Blue light photoreceptors in plants and animals. Annu Rev Plant Biol 62, 335–364. 10.1146/annurev-arplant-042110-103759.

16. Cashmore, A.R., Jarillo, J.A., Wu, Y.J., and Liu, D. (1999). Cryptochromes: Blue light receptors for plants and animals. Science (1979) 284, 760–765. 10.1126/science.284.5415.760.

17. Palayam, M., Ganapathy, J., Guercio, A.M., Tal, L., Deck, S.L., and Shabek, N. (2021). Structural insights into photoactivation of plant Cryptochrome-2. Commun Biol 4, 28. 10.1038/s42003-020-01531-x.

18. Yu, X., Sayegh, R., Maymon, M., Warpeha, K., Klejnot, J., Yang, H., Huang, J., Lee, J., Kaufman, L., and Lin, C. (2009). Formation of Nuclear Bodies of Arabidopsis CRY2 in Response to Blue Light Is Associated with Its Blue Light–Dependent Degradation. Plant Cell 21, 118–130. 10.1105/tpc.108.061663.

19. Liu, S., Zhang, L., Gao, L., Chen, Z., Bie, Y., Zhao, Q., Zhang, S., Hu, X., Liu, Q., Wang, X., et al. (2022). Differential photoregulation of the nuclear and cytoplasmic CRY1 in Arabidopsis. New Phytologist 234, 1332–1346. 10.1111/nph.18007.

20. Wang, X., Jiang, B., Gu, L., Chen, Y., Mora, M., Zhu, M., Noory, E., Wang, Q., and Lin, C. (2021). A photoregulatory mechanism of the circadian clock in Arabidopsis. Nat Plants 7, 1397–1408. 10.1038/s41477-021-01002-z.

21. Chen, Y., Hu, X., Liu, S., Su, T., Huang, H., Ren, H., Gao, Z., Wang, X., Lin, D., Wohlschlegel, J.A., et al. (2021). Regulation of Arabidopsis photoreceptor CRY2 by two distinct E3 ubiquitin ligases. Nat Commun 12, 2155. 10.1038/s41467-021-22410-x.

22. Liu, Q., Wang, Q., Liu, B., Wang, W., Wang, X., Park, J., Yang, Z., Du, X., Bian, M., and Lin, C. (2016). The Blue Light-Dependent Polyubiquitination and Degradation of Arabidopsis Cryptochrome2 Requires Multiple E3 Ubiquitin Ligases. Plant Cell Physiol 57, 2175–2186. 10.1093/pcp/pcw134.

23. Yu, X., Klejnot, J., Zhao, X., Shalitin, D., Maymon, M., Yang, H., Lee, J., Liu, X., Lopez, J., and Lin, C. (2007). Arabidopsis Cryptochrome 2 Completes Its Posttranslational Life Cycle in the Nucleus. Plant Cell 19, 3146–3156. 10.1105/tpc.107.053017.

24. Miao, L., Zhao, J., Yang, G., Xu, P., Cao, X., Du, S., Xu, F., Jiang, L., Zhang, S., Wei, X., et al. (2022). Arabidopsis cryptochrome 1 undergoes COP1 and LRBs-dependent degradation in response to high blue light. New Phytologist 234, 1347–1362. 10.1111/nph.17695.

25. Lindback, L.N., Hu, Y., Ackermann, A., Artz, O., and Pedmale, U. v. (2022). UBP12 and UBP13 deubiquitinases destabilize the CRY2 blue light receptor to regulate Arabidopsis growth. Current Biology 32, 1–11. 10.1016/j.cub.2022.05.046.

26. Wang, X., Wang, L., Huang, Y., Deng, Z., Li, C., Zhang, J., Zheng, M., and Yan, S. (2022). A plant-specific module for homologous recombination repair. Proceedings of the National Academy of Sciences 119. 10.1073/pnas.2202970119.

27. Brooks, C.L., and Gu, W. (2011). p53 regulation by ubiquitin. FEBS Lett 585, 2803–2809. 10.1016/j.febslet.2011.05.022.

28. Valles, G.J., Bezsonova, I., Woodgate, R., and Ashton, N.W. (2020). USP7 Is a Master Regulator of Genome Stability. Front Cell Dev Biol 8. 10.3389/fcell.2020.00717.

29. Sharma, N., Zhu, Q., Wani, G., He, J., Wang, Q.E., and Wani, A.A. (2014). USP3 counteracts RNF168 via deubiquitinating H2A and γh2AX at lysine 13 and 15. Cell Cycle 13, 106–114. 10.4161/cc.26814.

30. Hsu, D.S., Zhao, X., Zhao, S., Kazantsev, A., Wang, R.-P., Todo, T., Wei, Y.-F., and Sancar, A. (1996). Putative Human Blue-Light Photoreceptors hCRY1 and hCRY2 Are Flavoproteins. Biochemistry 35, 13871–13877. 10.1021/bi962209o.

31. Özgür, S., and Sancar, A. (2003). Purification and Properties of Human Blue-Light Photoreceptor Cryptochrome 2. Biochemistry 42, 2926–2932. 10.1021/bi026963n.

32. Papp, S.J., Huber, A.L., Jordan, S.D., Kriebs, A., Nguyen, M., Moresco, J.J., Yates, J.R., and Lamia, K.A. (2015). DNA damage shifts circadian clock time via Hausp-dependent Cry1 stabilization. Elife 4. 10.7554/eLife.04883.

33. Tissot, N., and Ulm, R. (2020). Cryptochrome-mediated blue-light signalling modulates UVR8 photoreceptor activity and contributes to UV-B tolerance in Arabidopsis. Nat Commun 11, 1323. 10.1038/s41467-020-15133-y.

34. Rai, N., O’Hara, A., Farkas, D., Safronov, O., Ratanasopa, K., Wang, F., Lindfors, A. v., Jenkins, G.I., Lehto, T., Salojärvi, J., et al. (2020). The photoreceptor UVR8 mediates the perception of both UV-B and UV-A wavelengths up to 350 nm of sunlight with responsivity moderated by cryptochromes. Plant Cell Environ 43, 1513–1527. 10.1111/pce.13752.

35. Rai, N., Neugart, S., Yan, Y., Wang, F., Siipola, S.M., Lindfors, A. v., Winkler, J.B., Albert, A., Brosché, M., Lehto, T., et al. (2019). How do cryptochromes and UVR8 interact in natural and simulated sunlight? J Exp Bot 70, 4975–4990. 10.1093/jxb/erz236.

36. Cui, X., Lu, F., Li, Y., Xue, Y., Kang, Y., Zhang, S., Qiu, Q., Zheng, X.C., Cui, X., Zheng, S., et al. (2013). Ubiquitin-specific proteases UBP12 and UBP13 act in circadian clock and photoperiodic flowering regulation in Arabidopsis. Plant Physiol 162, 897–906. 10.1104/pp.112.213009.

37. Molinier, J., Oakeley, E.J., Niederhauser, O., Kovalchuk, I., and Hohn, B. (2005). Dynamic response of plant genome to ultraviolet radiation and other genotoxic stresses. Mutation Research - Fundamental and Molecular Mechanisms of Mutagenesis 571, 235–247. 10.1016/j.mrfmmm.2004.09.016.

38. Takahashi, N., Inagaki, S., Nishimura, K., Sakakibara, H., Antoniadi, I., Karady, M., Ljung, K., and Umeda, M. (2021). Alterations in hormonal signals spatially coordinate distinct responses to DNA double-strand breaks in Arabidopsis roots. Sci Adv 7. 10.1126/sciadv.abg0993.

39. Tamura, K., Adachi, Y., Chiba, K., Oguchi, K., and Takahashi, H. (2002). Identification of Ku70 and Ku80 homologues in Arabidopsis thaliana: Evidence for a role in the repair of DNA double-strand breaks. Plant Journal 29, 771–781. 10.1046/j.1365-313X.2002.01258.x.

40. Castells, E., Molinier, J., Drevensek, S., Genschik, P., Barneche, F., and Bowler, C. (2010). det1-1-induced UV-C hyposensitivity through UVR3 and PHR1 photolyase gene over-expression. The Plant Journal 63, 392–404. 10.1111/j.1365-313X.2010.04249.x.

41. Ulm, R., Revenkova, E., di Sansebastiano, G.-P., Bechtold, N., and Paszkowski, J. (2001). Mitogen-activated protein kinase phosphatase is required for genotoxic stress relief in Arabidopsis. Genes Dev 15, 699–709. 10.1101/gad.192601.

42. Besteiro, M.A.G., Bartels, S., Albert, A., and Ulm, R. (2011). Arabidopsis MAP kinase phosphatase 1 and its target MAP kinases 3 and 6 antagonistically determine UV-B stress tolerance, independent of the UVR8 photoreceptor pathway. Plant Journal 68, 727–737. 10.1111/j.1365-313X.2011.04725.x.

43. Culligan, K.M., Robertson, C.E., Foreman, J., Doerner, P., and Britt, A.B. (2006). ATR and ATM play both distinct and additive roles in response to ionizing radiation. Plant Journal 48, 947–961. 10.1111/j.1365-313X.2006.02931.x.

44. Ogita, N., Okushima, Y., Tokizawa, M., Yamamoto, Y.Y., Tanaka, M., Seki, M., Makita, Y., Matsui, M., Okamoto-Yoshiyama, K., Sakamoto, T., et al. (2018). Identifying the target genes of SUPPRESSOR OF GAMMA RESPONSE 1, a master transcription factor controlling DNA damage response in Arabidopsis. Plant Journal 94, 439–453. 10.1111/tpj.13866.

45. Li, W., Chen, C., Markmann-Mulisch, U., Timofejeva, L., Schmelzer, E., Ma, H., and Reiss, B. (2004). The Arabidopsis AtRAD51 gene is dispensable for vegetative development but required for meiosis. Proceedings of the National Academy of Sciences 101, 10596–10601. 10.1073/pnas.0404110101.

46. Ulm, R., Baumann, A., Oravecz, A., Máté, Z., Ádám, É., Oakeley, E.J., Schäfer, E., and Nagy, F. (2004). Genome-wide analysis of gene expression reveals function of the bZIP transcription factor HY5 in the UV-B response of Arabidopsis. Proceedings of the National Academy of Sciences 101, 1397–1402. 10.1073/pnas.0308044100.

47. Rodriguez, E., Chevalier, J., el Ghoul, H., Voldum-Clausen, K., Mundy, J., and Petersen, M. (2018). DNA damage as a consequence of NLR activation. PLoS Genet 14, e1007235. 10.1371/journal.pgen.1007235.

48. Jin, H. (2000). Transcriptional repression by AtMYB4 controls production of UV-protecting sunscreens in Arabidopsis. EMBO J 19, 6150–6161. 10.1093/emboj/19.22.6150.

49. Kliebenstein, D.J., Lim, J.E., Landry, L.G., and Last, R.L. (2002). Arabidopsis UVR8 regulates ultraviolet-B signal transduction and tolerance and contains sequence similarity to human Regulator of Chromatin Condensation 1. Plant Physiol 130, 234–243. 10.1104/pp.005041.

50. Tsurumoto, T., Fujikawa, Y., Onoda, Y., Ochi, Y., Ohta, D., and Okazawa, A. (2022). Transcriptome and metabolome analyses revealed that narrowband 280 and 310 nm UV-B induce distinctive responses in Arabidopsis. Sci Rep 12, 4319. 10.1038/s41598-022-08331-9.

51. Danon, A., Rotari, V.I., Gordon, A., Mailhac, N., and Gallois, P. (2004). Ultraviolet-C Overexposure Induces Programmed Cell Death in Arabidopsis, Which Is Mediated by Caspase-like Activities and Which Can Be Suppressed by Caspase Inhibitors, p35 and Defender against Apoptotic Death. Journal of Biological Chemistry 279, 779–787. 10.1074/jbc.M304468200.

52. Ernst, J., Vainas, O., Harbison, C.T., Simon, I., and Bar-Joseph, Z. (2007). Reconstructing dynamic regulatory maps. Mol Syst Biol 3, 74. 10.1038/msb4100115.

53. Schulz, M.H., Devanny, W.E., Gitter, A., Zhong, S., Ernst, J., and Bar-Joseph, Z. (2012). DREM 2.0: Improved reconstruction of dynamic regulatory networks from time-series expression data. BMC Syst Biol 6, 104. 10.1186/1752-0509-6-104.

54. Heinz, S., Benner, C., Spann, N., Bertolino, E., Lin, Y.C., Laslo, P., Cheng, J.X., Murre, C., Singh, H., and Glass, C.K. (2010). Simple Combinations of Lineage-Determining Transcription Factors Prime cis-Regulatory Elements Required for Macrophage and B Cell Identities. Mol Cell 38, 576–589. 10.1016/j.molcel.2010.05.004.

55. Benn, G., Wang, C.-Q., Hicks, D.R., Stein, J., Guthrie, C., and Dehesh, K. (2014). A key general stress response motif is regulated non-uniformly by CAMTA transcription factors. The Plant Journal 80, 82–92. 10.1111/tpj.12620.

56. Iqbal, Z., Shariq Iqbal, M., Singh, S.P., and Buaboocha, T. (2020). Ca2+/Calmodulin Complex Triggers CAMTA Transcriptional Machinery Under Stress in Plants: Signaling Cascade and Molecular Regulation. Front Plant Sci 11. 10.3389/fpls.2020.598327.

57. Matsumura, M., Nomoto, M., Itaya, T., Aratani, Y., Iwamoto, M., Matsuura, T., Hayashi, Y., Mori, T., Skelly, M.J., Yamamoto, Y.Y., et al. (2022). Mechanosensory trichome cells evoke a mechanical stimuli–induced immune response in Arabidopsis thaliana. Nat Commun 13, 1216. 10.1038/s41467-022-28813-8.

58. Thompson, M. v., and Wolniak, S.M. (2008). A Plasma Membrane-Anchored Fluorescent Protein Fusion Illuminates Sieve Element Plasma Membranes in Arabidopsis and Tobacco. Plant Physiol 146, 1599–1610. 10.1104/pp.107.113274.

59. Banerjee, R., Schleicher, E., Meier, S., Viana, R.M., Pokorny, R., Ahmad, M., Bittl, R., and Batschauer, A. (2007). The Signaling State of Arabidopsis Cryptochrome 2 Contains Flavin Semiquinone. Journal of Biological Chemistry 282, 14916–14922. 10.1074/jbc.M700616200.

60. Che, D.L., Duan, L., Zhang, K., and Cui, B. (2015). The Dual Characteristics of Light-Induced Cryptochrome 2, Homo-oligomerization and Heterodimerization, for Optogenetic Manipulation in Mammalian Cells. ACS Synth Biol 4, 1124–1135. 10.1021/acssynbio.5b00048.

61. Liu, H., Yu, X., Li, K., Klejnot, J., Yang, H., Lisiero, D., and Lin, C. (2008). Photoexcited CRY2 interacts with CIB1 to regulate transcription and floral initiation in Arabidopsis. Science (1979) 322, 1535–1539. 10.1126/science.1163927.

62. Shafi, A.A., McNair, C.M., McCann, J.J., Alshalalfa, M., Shostak, A., Severson, T.M., Zhu, Y., Bergman, A., Gordon, N., Mandigo, A.C., et al. (2021). The circadian cryptochrome, CRY1, is a pro-tumorigenic factor that rhythmically modulates DNA repair. Nat Commun 12, 401. 10.1038/s41467-020-20513-5.

63. Jiang, C.-Z., Yee, J., Mitchell, D.L., and Britt, A.B. (1997). Photorepair mutants of Arabidopsis. Proceedings of the National Academy of Sciences 94, 7441–7445. 10.1073/pnas.94.14.7441.

64. Molinier, J., Lechner, E., Dumbliauskas, E., and Genschik, P. (2008). Regulation and role of arabidopsis CUL4-DDB1A-DDB2 in maintaining genome integrity upon UV stress. PLoS Genet 4. 10.1371/journal.pgen.1000093.

65. Lee, J., He, K., Stolc, V., Lee, H., Figueroa, P., Gao, Y., Tongprasit, W., Zhao, H., Lee, I., and Xing, W.D. (2007). Analysis of transcription factor HY5 genomic binding sites revealed its hierarchical role in light regulation of development. Plant Cell 19, 731–749. 10.1105/tpc.106.047688.

66. Ponnu, J., Riedel, T., Penner, E., Schrader, A., and Hoecker, U. (2019). Cryptochrome 2 competes with COP1 substrates to repress COP1 ubiquitin ligase activity during Arabidopsis photomorphogenesis. Proc Natl Acad Sci U S A 116, 27133–27141. 10.1073/pnas.1909181116.

67. Lau, K., Podolec, R., Chappuis, R., Ulm, R., and Hothorn, M. (2019). Plant photoreceptors and their signaling components compete for COP 1 binding via VP peptide motifs. EMBO J 38. 10.15252/embj.2019102140.

68. Castells, E., Molinier, J., Benvenuto, G., Bourbousse, C., Zabulon, G., Zalc, A., Cazzaniga, S., Genschik, P., Barneche, F., and Bowler, C. (2011). The conserved factor DE-ETIOLATED 1 cooperates with CUL4-DDB1 DDB2 to maintain genome integrity upon UV stress. EMBO J 30, 1162–1172. 10.1038/emboj.2011.20.

69. Oberle, C., and Blattner, C. (2010). Regulation of the DNA Damage Response to DSBs by Post-Translational Modifications. Curr Genomics 11, 184–198. 10.2174/138920210791110979.

70. Gross, S., Rahal, R., Stransky, N., Lengauer, C., and Hoeflich, K.P. (2015). Targeting cancer with kinase inhibitors. Journal of Clinical Investigation 125, 1780–1789. 10.1172/JCI76094.

71. Hafner, A., Bulyk, M.L., Jambhekar, A., and Lahav, G. (2019). The multiple mechanisms that regulate p53 activity and cell fate. Nat Rev Mol Cell Biol 20, 199–210. 10.1038/s41580-019-0110-x.

72. Doil, C., Mailand, N., Bekker-Jensen, S., Menard, P., Larsen, D.H., Pepperkok, R., Ellenberg, J., Panier, S., Durocher, D., Bartek, J., et al. (2009). RNF168 Binds and Amplifies Ubiquitin Conjugates on Damaged Chromosomes to Allow Accumulation of Repair Proteins. Cell 136, 435–446. 10.1016/j.cell.2008.12.041.

73. Li, M., Chen, D., Shiloh, A., Luo, J., Nikolaev, A.Y., Qin, J., and Gu, W. (2002). Deubiquitination of p53 by HAUSP is an important pathway for p53 stabilization. Nature 416, 648–653. 10.1038/nature737.

74. Alonso-de Vega, I., Martín, Y., and Smits, V.A. (2014). USP7 controls Chk1 protein stability by direct deubiquitination. Cell Cycle 13, 3921–3926. 10.4161/15384101.2014.973324.

75. Sanchez, Y., Wong, C., Thoma, R.S., Richman, R., Wu, Z., Piwnica-Worms, H., and Elledge, S.J. (1997). Conservation of the Chk1 Checkpoint Pathway in Mammals: Linkage of DNA Damage to Cdk Regulation Through Cdc25. Science (1979) 277, 1497–1501. 10.1126/science.277.5331.1497.

76. al Khateeb, W.M., Sher, A.A., Marcus, J.M., and Schroeder, D.F. (2019). UVSSA, UBP12, and RDO2/TFIIS Contribute to Arabidopsis UV Tolerance. Front Plant Sci 10. 10.3389/fpls.2019.00516.

77. Derkacheva, M., Liu, S., Figueiredo, D.D., Gentry, M., Mozgova, I., Nanni, P., Tang, M., Mannervik, M., Köhler, C., and Hennig, L. (2016). H2A deubiquitinases UBP12/13 are part of the Arabidopsis polycomb group protein system. Nat Plants 2, 16126. 10.1038/nplants.2016.126.

78. Mattiroli, F., and Penengo, L. (2021). Histone Ubiquitination: An Integrative Signaling Platform in Genome Stability. Trends in Genetics 37, 566–581. 10.1016/j.tig.2020.12.005.

79. Jackson, S.P., and Bartek, J. (2009). The DNA-damage response in human biology and disease. Nature 461, 1071–1078. 10.1038/nature08467.

80. Jones, S.N., Roe, A.E., Donehower, L.A., and Bradley, A. (1995). Rescue of embryonic lethality in Mdm2-deficient mice by absence of p53. Nature 378, 206–208. 10.1038/378206a0.

81. Bouché, N., Scharlat, A., Snedden, W., Bouchez, D., and Fromm, H. (2002). A Novel Family of Calmodulin-binding Transcription Activators in Multicellular Organisms. Journal of Biological Chemistry 277, 21851–21861. 10.1074/jbc.M200268200.

82. Song, K., Backs, J., McAnally, J., Qi, X., Gerard, R.D., Richardson, J.A., Hill, J.A., Bassel-Duby, R., and Olson, E.N. (2006). The Transcriptional Coactivator CAMTA2 Stimulates Cardiac Growth by Opposing Class II Histone Deacetylases. Cell 125, 453–466. 10.1016/j.cell.2006.02.048.

83. Long, C., Grueter, C.E., Song, K., Qin, S., Qi, X., Kong, Y.M., Shelton, J.M., Richardson, J.A., Zhang, C.-L., Bassel-Duby, R., et al. (2014). Ataxia and Purkinje cell degeneration in mice lacking the CAMTA1 transcription factor. Proceedings of the National Academy of Sciences 111, 11521–11526. 10.1073/pnas.1411251111.

84. Bas-Orth, C., Tan, Y.-W., Oliveira, A.M.M., Bengtson, C.P., and Bading, H. (2016). The calmodulin-binding transcription activator CAMTA1 is required for long-term memory formation in mice. Learning & Memory 23, 313–321. 10.1101/lm.041111.115.

85. Schraivogel, D., Weinmann, L., Beier, D., Tabatabai, G., Eichner, A., Zhu, J.Y., Anton, M., Sixt, M., Weller, M., Beier, C.P., et al. (2011). CAMTA1 is a novel tumour suppressor regulated by miR-9/9 * in glioblastoma stem cells. EMBO J 30, 4309–4322. 10.1038/emboj.2011.301.

86. Yoshiyama, K., Conklin, P.A., Huefner, N.D., and Britt, A.B. (2009). Suppressor of gamma response 1 (SOG1) encodes a putative transcription factor governing multiple responses to DNA damage. Proc Natl Acad Sci U S A 106, 12843–12848. 10.1073/pnas.0810304106.

87. Winter, D., Vinegar, B., Nahal, H., Ammar, R., Wilson, G. v., and Provart, N.J. (2007). An “Electronic Fluorescent Pictograph” Browser for Exploring and Analyzing Large-Scale Biological Data Sets. PLoS One 2, e718. 10.1371/journal.pone.0000718.

88. Horvath, B.M., Kourova, H., Nagy, S., Nemeth, E., Magyar, Z., Papdi, C., Ahmad, Z., Sanchez-Perez, G.F., Perilli, S., Blilou, I., et al. (2017). Arabidopsis RETINOBLASTOMA RELATED directly regulates DNA damage responses through functions beyond cell cycle control. EMBO J 36, 1261–1278. 10.15252/embj.201694561.

89. Li, S., Lavagnino, Z., Lemacon, D., Kong, L., Ustione, A., Ng, X., Zhang, Y., Wang, Y., Zheng, B., Piwnica-Worms, H., et al. (2019). Ca2+-Stimulated AMPK-Dependent Phosphorylation of Exo1 Protects Stressed Replication Forks from Aberrant Resection. Mol Cell 74, 1123–1137.e6. 10.1016/j.molcel.2019.04.003.

90. Guo, H., Yang, H., Mockler, T.C., and Lin, C. (1998). Regulation of flowering time by Arabidopsis photoreceptors. Science (1979) 279, 1360–1363. 10.1126/science.279.5355.1360.

91. Lin, C., Yang, H., Guo, H., Mockler, T., Chen, J., and Cashmore, A.R. (1998). Enhancement of blue-light sensitivity of Arabidopsis seedlings by a blue light receptor cryptochrome 2. Proc Natl Acad Sci U S A 95, 2686–2690. 10.1073/pnas.95.5.2686.

92. Yagi, K., Ozawa, T., and Harada, M. (1959). Change of Absorption Spectrum of Flavin Adenine Dinucleotide byits Binding with both D-Amino Acid Oxidase Apo-Protein and Benzoate. Nature 184, 1938–1939. 10.1038/1841938a0.

93. Zeng, Z., Wei, J., Liu, Y., Zhang, W., and Mabe, T. (2018). Magnetoreception of Photoactivated Cryptochrome 1 in Electrochemistry and Electron Transfer. ACS Omega 3, 4752–4759. 10.1021/acsomega.8b00645.

94. Mockler, T.C., Guo, H., Yang, H., Duong, H., and Lin, C. (1999). Antagonistic actions of Arabidopsis cryptochromes and phytochrome B in the regulation of floral induction. Development 126, 2073–2082. 10.1242/dev.126.10.2073.

95. Kannan, K., Nelson, A.D.L., and Shippen, D.E. (2008). Dyskerin Is a Component of the Arabidopsis Telomerase RNP Required for Telomere Maintenance. Mol Cell Biol 28, 2332–2341. 10.1128/MCB.01490-07.

96. Deng, X.W., Matsui, M., Wei, N., Wagner, D., Chu, A.M., Feldmann, K.A., and Quail, P.H. (1992). COP1, an arabidopsis regulatory gene, encodes a protein with both a zinc-binding motif and a Gβ homologous domain. Cell 71, 791–801. 10.1016/0092-8674(92)90555-Q.

97. Kim, Y., Gilmour, S.J., Chao, L., Park, S., and Thomashow, M.F. (2020). Arabidopsis CAMTA Transcription Factors Regulate Pipecolic Acid Biosynthesis and Priming of Immunity Genes. Mol Plant 13, 157–168. 10.1016/j.molp.2019.11.001.

98. Pedmale, U. v., Huang, S.S.C., Zander, M., Cole, B.J., Hetzel, J., Ljung, K., Reis, P.A.B., Sridevi, P., Nito, K., Nery, J.R., et al. (2016). Cryptochromes Interact Directly with PIFs to Control Plant Growth in Limiting Blue Light. Cell 164, 233–245. 10.1016/j.cell.2015.12.018.

99. Karimi, M., Bleys, A., Vanderhaeghen, R., and Hilson, P. (2007). Building Blocks for Plant Gene Assembly. Plant Physiol 145, 1183–1191. 10.1104/pp.107.110411.

100. Clough, S.J., and Bent, A.F. (1998). Floral dip: a simplified method forAgrobacterium-mediated transformation ofArabidopsis thaliana. The Plant Journal 16, 735–743. 10.1046/j.1365-313x.1998.00343.x.

101. Porebski, S., Bailey, L.G., and Baum, B.R. (1997). Modification of a CTAB DNA extraction protocol for plants containing high polysaccharide and polyphenol components. Plant Mol Biol Report 15, 8–15. 10.1007/BF02772108.

102. Schneider, C.A., Rasband, W.S., and Eliceiri, K.W. (2012). NIH Image to ImageJ: 25 years of image analysis. Nat Methods 9, 671–675. 10.1038/nmeth.2089.

103. Dobin, A., Davis, C.A., Schlesinger, F., Drenkow, J., Zaleski, C., Jha, S., Batut, P., Chaisson, M., and Gingeras, T.R. (2013). STAR: Ultrafast universal RNA-seq aligner. Bioinformatics 29, 15–21. 10.1093/bioinformatics/bts635.

104. Trapnell, C., Roberts, A., Goff, L., Pertea, G., Kim, D., Kelley, D.R., Pimentel, H., Salzberg, S.L., Rinn, J.L., and Pachter, L. (2012). Differential gene and transcript expression analysis of RNA-seq experiments with TopHat and Cufflinks. Nat Protoc 7, 562–578. 10.1038/nprot.2012.016.

105. Love, M.I., Huber, W., and Anders, S. (2014). Moderated estimation of fold change and dispersion for RNA-seq data with DESeq2. Genome Biol 15, 550. 10.1186/s13059-014-0550-8.

106. Mi, H., and Thomas, P. (2009). PANTHER Pathway: An Ontology-Based Pathway Database Coupled with Data Analysis Tools. In, pp. 123–140. 10.1007/978-1-60761-175-2_7.

107. Livak, K.J., and Schmittgen, T.D. (2001). Analysis of Relative Gene Expression Data Using Real-Time Quantitative PCR and the 2-ΔΔCT Method. Methods 25, 402–408. 10.1006/meth.2001.1262.

