## Supplementary figures and images for "Cryptochromes and UBP12/13 deubiquitinases antagonistically regulate DNA damage response in Arabidopsis"

### Figure S1

**Figure S1, Hu et al.**

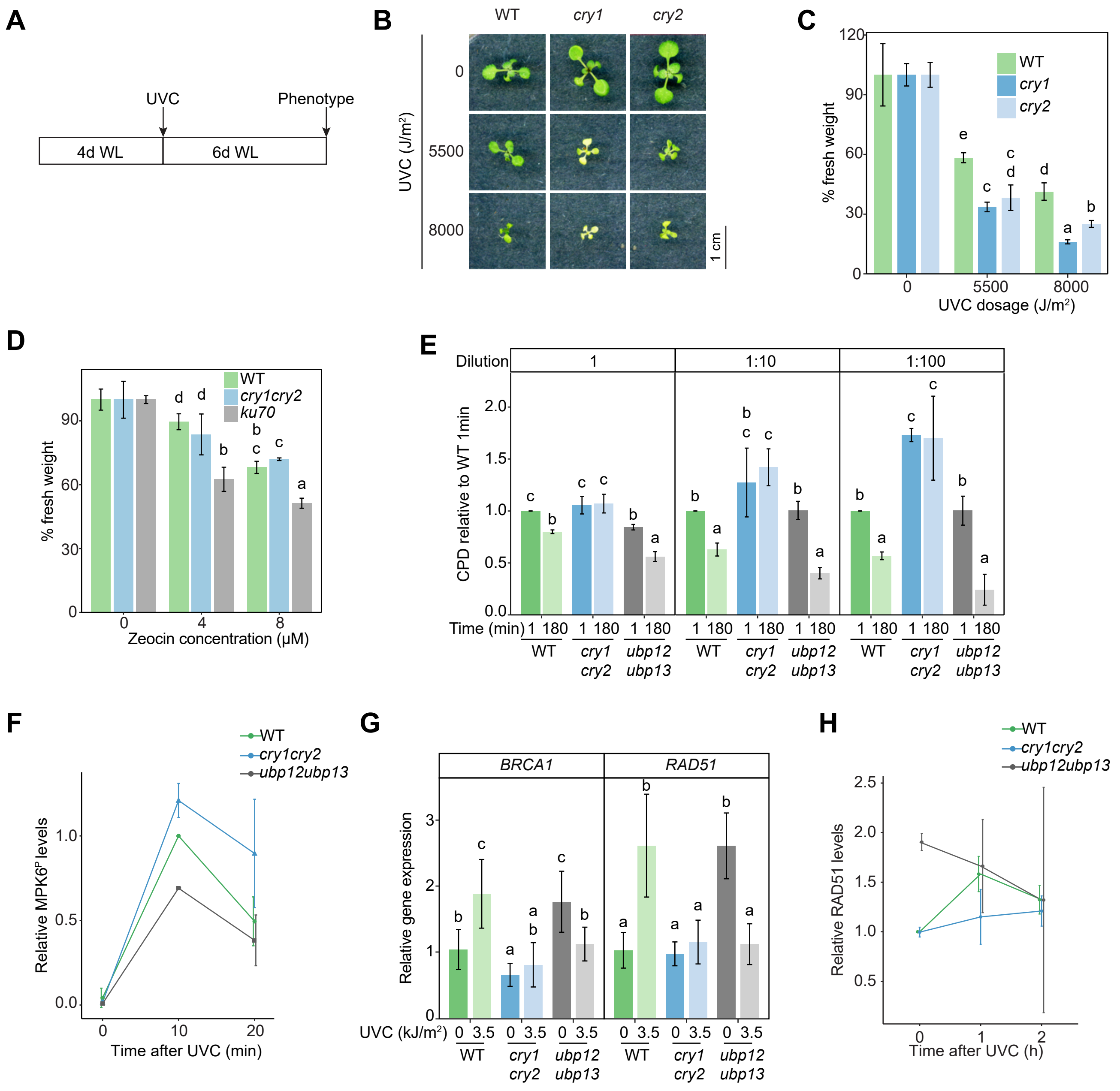

### Figure S2

**Figure S2, Hu et al.**

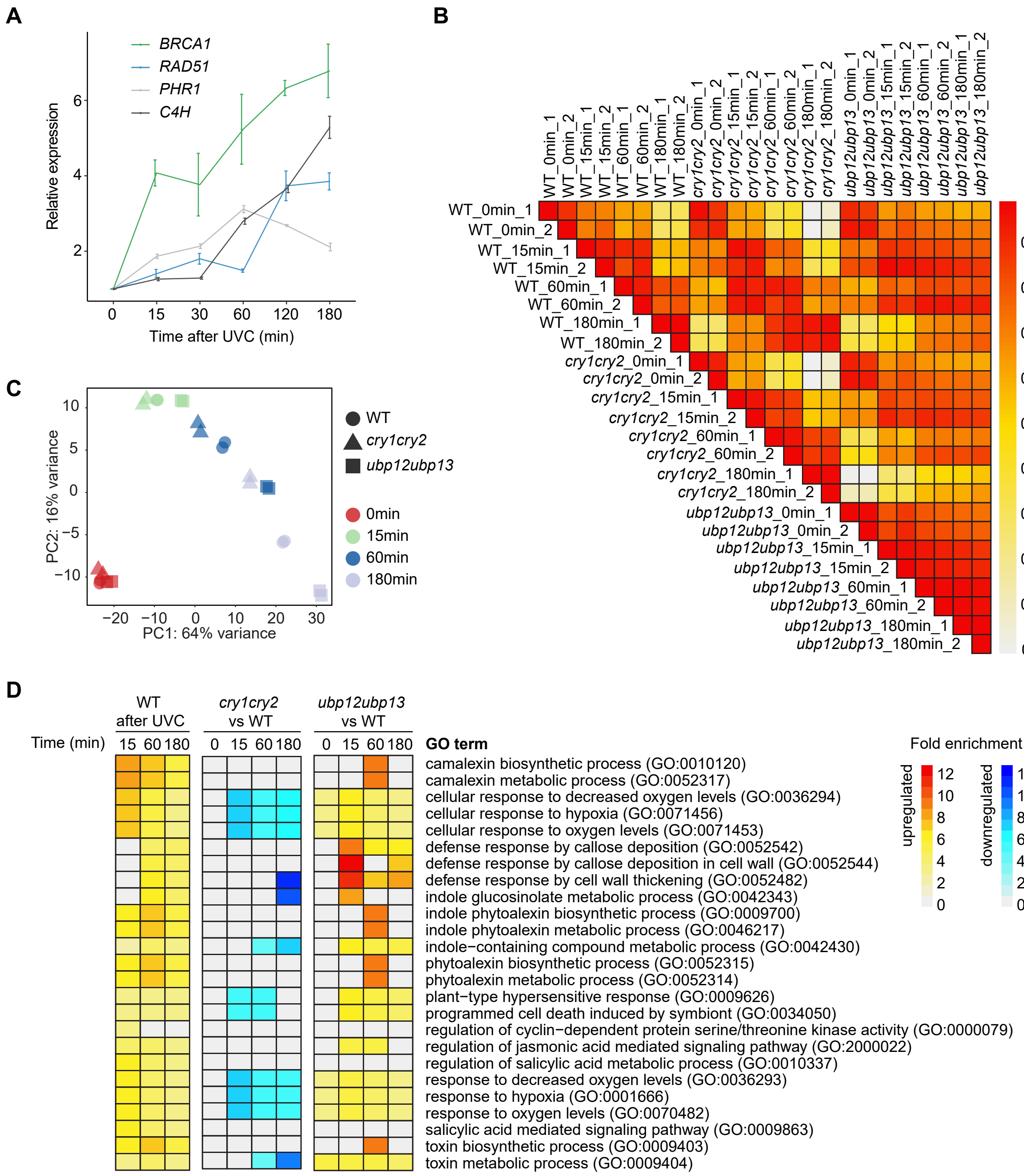

### Figure S4

#### Figure S4, Hu et al.

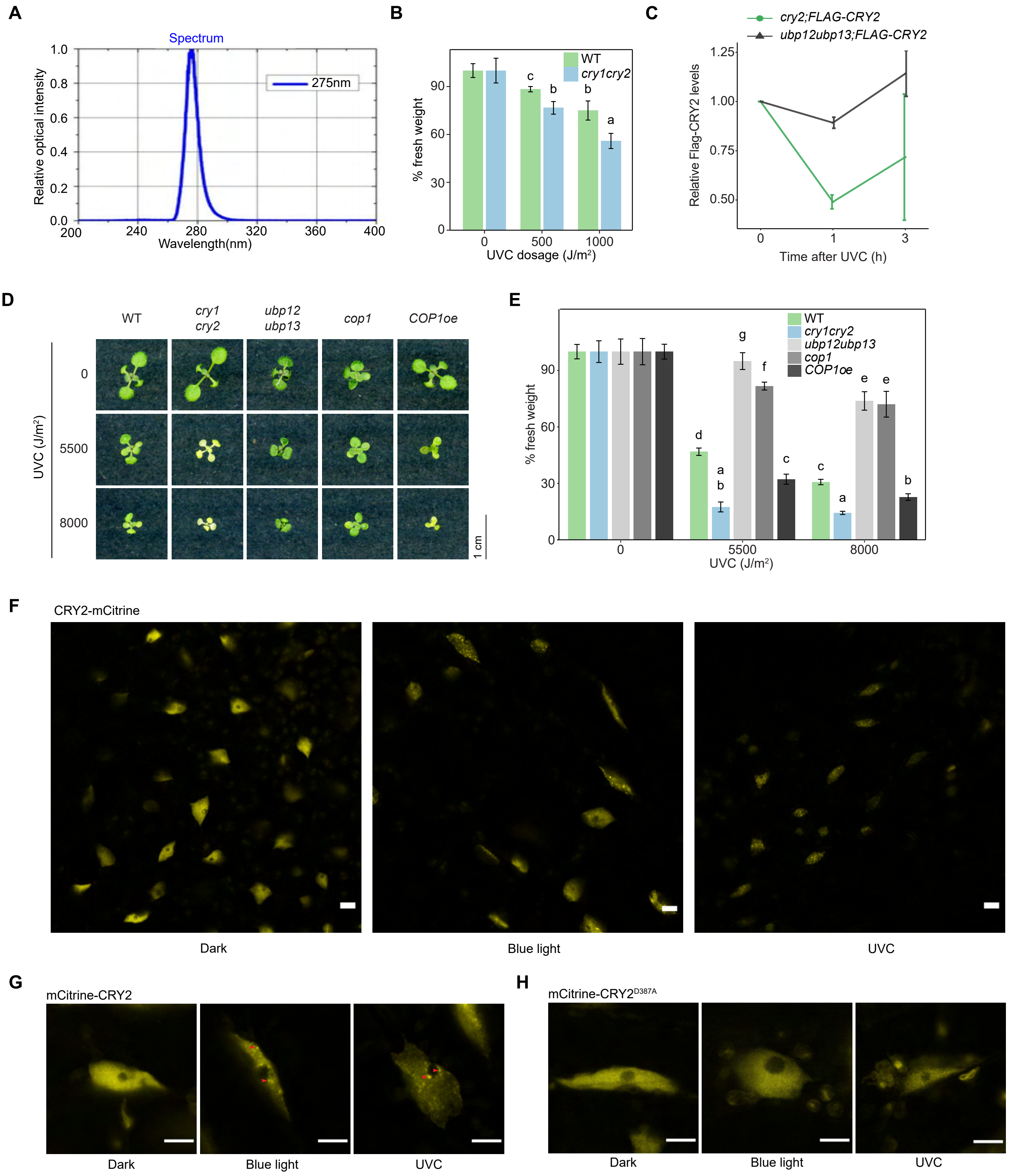
