## Supplementary material for "Cryptochromes and UBP12/13 deubiquitinases antagonistically regulate DNA damage response in Arabidopsis": Figure S3

**Figure S3, Hu et al.**

**A**

Motifs identified in promoters of upregulated genes

| Motif | *P* value | Best match |
|---|---|---|
| 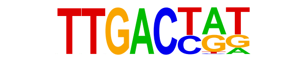 | $10^{-26}$ | WRKY18(WRKY) |
| 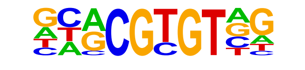 | $10^{-25}$ | **CMTA2** |
| 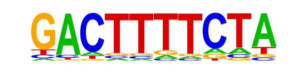 | $10^{-16}$ | WRKY50 |
| 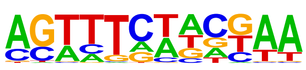 | $10^{-16}$ | At1g25550(G2like) |
| 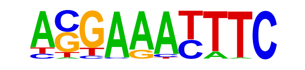 | $10^{-15}$ | EPR1(MYBrelated) |
| 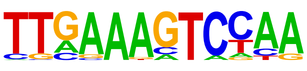 | $10^{-15}$ | WRKY50 |
| 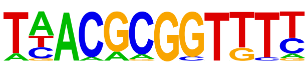 | $10^{-13}$ | **CAMTA1** |
| 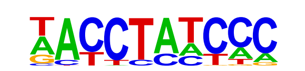 | $10^{-12}$ | MYB3 |
| 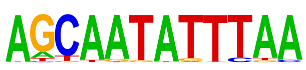 | $10^{-12}$ | HAT5 |
| 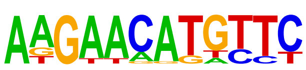 | $10^{-12}$ | KAN2 |

**B**

Motifs identified in promoters of downregulated genes

| Motif | *P* value | Best match |
|---|---|---|
| 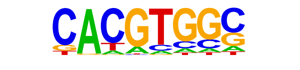 | $10^{-22}$ | PIF4 |
| 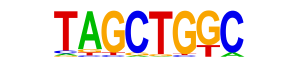 | $10^{-19}$ | bZIP18(bZIP) |
| 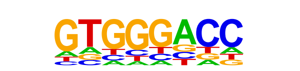 | $10^{-18}$ | TCP20(TCP) |
| 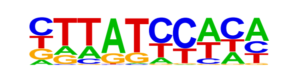 | $10^{-16}$ | At1g1900(MYBrelated) |
| 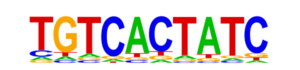 | $10^{-14}$ | At5g04390(C2H2) |
| 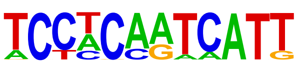 | $10^{-14}$ | ATHB7(Homeobox) |
| 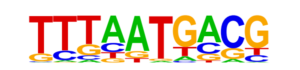 | $10^{-14}$ | At1g76110(ARID) |
| 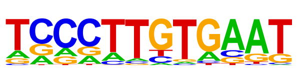 | $10^{-13}$ | AT2G20110 |
| 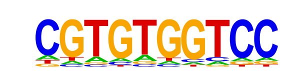 | $10^{-12}$ | TCP3(TCP) |
| 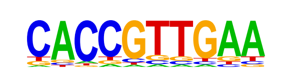 | $10^{-12}$ | MYB3R5(MYB) |

**C**

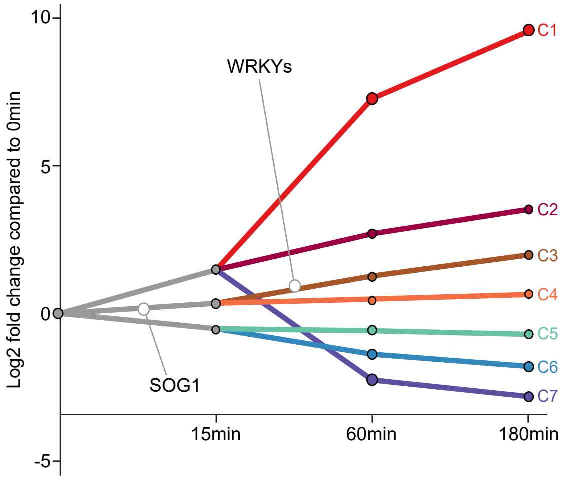


*cry1cry2*

**D**

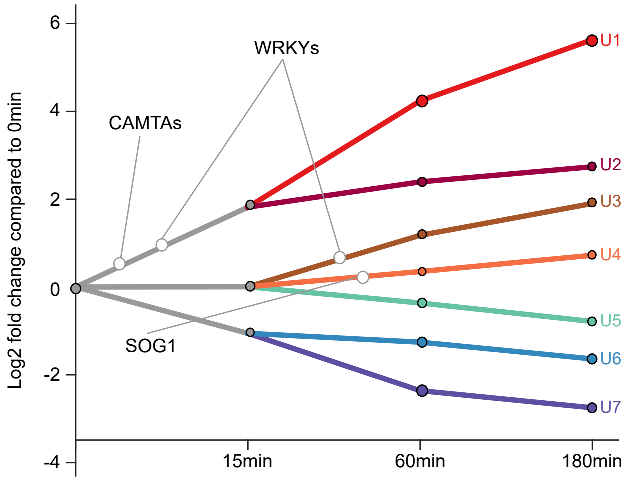


*ubp12ubp13*
