## Supplemental figure legends for "Cryptochromes and UBP12/13 deubiquitinases antagonistically regulate DNA damage response in Arabidopsis"

**Figure S1.**

(A) Schematic diagram illustrating the UVC treatment and phenotyping. 4-day-old seedlings grown in continuous white light (WL) were treated with UVC and grown in continuous WL for another 6 d before examination of phenotype.

(B) Phenotype of representative 10-day-old seedlings of the indicated genotypes treated with indicated UVC doses as described in (A).

(C) Fresh weight of 10-day-old seedlings of the indicated genotypes treated with indicated UVC doses as described in (A). Fresh weight was normalized to the untreated (0 $J/m^2$) samples of the same genotype, n = 3.

(D) Fresh weight of 12-day-old seedlings of the indicated genotypes treated with indicated concentrations of zeocin. 4-day-old seedlings grown in light and regular growth medium were transferred to a growth medium containing zeocin and grown for another 8 d before examination of phenotype. Fresh weight was normalized to the untreated (0 μM) samples of the same genotype. n = 3.

(E) Quantification of relative CPD levels using replicates of the dot blot shown in Figure 1D. The CPD levels were normalized to the average levels in WT at 1 min after UVC exposure for each dilution. One-way ANOVA analysis was separately performed for each dilution. n = 3.

(F) Quantification of phosphorylated MPK6 ($MPK6^P$) levels from replicates of Figure S1I. $MPK6^P$ levels were normalized to WT at 10 min for each blot. n = 2.

(G) Relative expression of *BRCA1* and *RAD51* genes derived from RT-qPCR analysis. 3-day-old etiolated seedlings were untreated (0 $kJ/m^2$) or treated with 3.5 $kJ/m^2$ of UVC and incubated in white light for 1.5 h before tissue collection. n = 3.

(H) Quantification of RAD51 protein levels using replicates of Figure S1G. RAD51 levels were normalized to WT at 0 h for each blot. n = 2.

For (C), (D), (E) and (G), different letters mean $p<0.05$ for one-way ANOVA analysis followed by Fisher's LSD posthoc test.

For (C-H), data show means ± standard deviation (SD) of independent replicates.

**Figure S2.**

(A) Relative expression of UV-responsive genes at indicated time points after UVC treatment derived from qRT-PCR analysis. 5-day-old light-grown WT seedlings were treated with 6000 J/m$^2$ of UVC and collected after 0, 15, 30, 60, 120, and 180 min. Data show means ± SD. n = 3.

(B) Pearson's correlation coefficient between all 24 RNA-seq samples. Coefficient was calculated based on the FPKM values of all genes in the transcriptome.

(C) Principal component analysis of all RNA-seq samples using normalized read counts.

(D) Heatmap of the top 10 GO terms enriched in the upregulated genes of WT at 15, 60 and 180 min.

**Figure S3.**

(A, B) Top 10 significant *cis*-regulatory motifs identified in the promoters of all upregulated (A) and downregulated (B) genes in WT after UVC.

(C, D) DREM model showing co-expressed gene groups and predicted transcription factors of all DEGs in *cry1cry2* (C) and *ubp12ubp13* (D) after UVC.

**Figure S4.**

(A) Spectrum of the light emitted by the UVC LED source adapted from the data sheet provided by the manufacturer (International Light Technologies, 2022).

(B) Fresh weight of 10-day-old seedlings with indicated genotypes treated with indicated UVC doses by the UVC LED source. 5-day-old light-grown seedlings were treated with UVC and incubated in light for another 5 days before examination of phenotype. Fresh weight was normalized to the untreated (0 J/m$^2$) samples of the same genotype. n = 3.

(C) Quantification of FLAG-CRY2 levels using replicates of Figure 4B. FLAG-CRY2 levels were normalized to the 0 h sample of the same genetic background. Data show means ± SD, n = 2.

(D) Phenotype of representative 10-day-old seedlings of the indicated genotypes treated with indicated UVC doses. 4-day-old light-grown seedlings were treated with UVC and then returned to white light for 6 days before examination of the phenotype.

(E) Fresh weight of 10-day-old seedlings of the indicated genotypes treated as in (D). The fresh weight percentage was calculated as described in (C). n = 3.

(F- H) Representative confocal microscopy images of transgenic *Arabidopsis* seedlings expressing CRY2 and mCitrine fusions. Seedlings were dark grown and either kept in dark or treated with blue light or UVC and fixed before imaging. C-terminal CRY2-mCitrine fusion in (F), and N-terminal mCitrine-CRY2 and mCitrine-CRY2$^{D387A}$ in (G) and (H), respectively.

For (B) and (E), different letters mean $p<0.05$ for one-way ANOVA analysis followed by Fisher's LSD posthoc test. Data show means ± SD.

For (F-H), the scale bar is 5 μm.

**Reference**

International Light Technologies (2022). UVC LED Module Array Data Sheet.
